# Pupil size and pupil change correlate with distinct cortical neurochemical systems across the adult lifespan

**DOI:** 10.64898/2026.08.24.746786

**Authors:** Elizabeth Riley, Eve De Rosa, Adam Anderson

## Abstract

Pupillometry is often used as a proxy for locus coeruleus (LC) activity but whether this relationship changes across the lifespan is unknown. We collected simultaneous pupillometry and multi-echo fMRI in 89 adults aged 19-78 during a visual oddball task. Oddball-evoked pupillary responses were present in all age groups and largest in middle age. Oddball-evoked LC BOLD responses were detected only in younger adults and did not predict individual pupillary responses. Pupil size and its first derivative (pupil change) correlated with whole-brain BOLD on distinct timescales and in distinct spatial patterns. Against 19 PET maps, pupil derivative coupling correlated uniquely with norepinephrine transporter (NET) density, and pupil size-unique coupling with μ-opioid, cannabinoid CB1, and glutamatergic mGluR5 density. Only the pupil derivative-NET relationship weakened with age, though it remained significant in every group. Pupil size and pupil change therefore carry different information, and only pupil change correlates with activity in NET-rich regions across the lifespan.

## Introduction

Pupil size changes to adapt to light conditions, but when light conditions are steady, pupil size changes size according to central nervous system control. In the absence of changes in ambient light, pupil size can be modulated by mental states and associated brain activity (Hartmann & Fischer, 2014), providing insights into cognitive phenomena such as affect (Kinner et al., 2017), attentional engagement (Unsworth & Robison, 2016), cognitive load (Granholm et al., 2017; Kahneman & Beatty, 1966) , and orienting responses (Riley et al., 2023). In addition, pupillometry has been applied to evaluate neurocognitive aging in older adults (El Haj et al., 2024; Elman et al., 2017; Frost et al., 2017; Hämmerer et al., 2017; He et al., 2020). Both tonic pupil size (Kuwamizu et al., 2022) and phasic pupillary responses (for e.g., He et al., 2020; Riley et al., 2023) have been correlated with various neural and cognitive processes. These investigations aim to make use of pupillometry as an accessible and cost-effective means of assessing brain health and cognition. In particular, the cognitive regulation of the pupil has been employed as a direct read out of the activity in the locus coeruleus (LC) of the brainstem (Costa & Rudebeck, 2016; Elman et al., 2017; Gilzenrat et al., 2010; Joshi et al., 2016; Murphy et al., 2014) - a neuromodulatory nucleus known for its role in regulating orienting (Aston-Jones & Cohen, 2005; Sara & Bouret, 2012), arousal (see Ross & Van Bockstaele, 2021), and memory (see Poe et al., 2020) and as the brain’s primary source of norepinephrine (NE, Schwarz & Luo, 2015). It is also the first place in the brain to accumulate Alzheimer’s-related tau pathology (Arnsten et al., 2021; Braak et al., 2011; Bueichekú et al., 2024), both underlining its importance as a topic of study. However, for exactly the reason that the LC is so important to study in middle-aged and older adults, it may also be unreliable, since it is unclear whether the known links between pupil size and LC activity are robust to pathological change in the LC. In order to be able to capitalize on pupil measurements as a window into the aging brain, it is necessary to conduct a robust characterization of pupillary dynamics and their degree of association with candidate neurotransmitter systems in adults across the lifespan.

Non-human animal research has demonstrated that rapid changes in pupil size are tightly coupled with neuronal activity in the LC (Grujic et al., 2024), while slow, low frequency change is more associated with general arousal level (Reimer et al., 2016). Similar associations have been observed in humans, with changes in pupil size serving as an indirect marker of LC activity in certain contexts (Costa & Rudebeck, 2016; Joshi et al., 2016; Murphy et al., 2014).

However, this association does not hold across all contexts (Grujic et al., 2024). The extent to which pupil metrics reliably reflect LC activity, or activity in any other particular brain region or system, may depend on task demands or experimental conditions. Depending on context, pupillary responses may represent NE, but also acetylcholine (Fotiou et al., 2009), dopamine (Bartošová et al., 2018) or serotonin signaling (Noehr-Jensen et al., 2009), or a combination (Larsen & Waters, 2018). This variability raises serious concerns about using pupillometry as a direct readout of LC function or NE release. To specifically assess LC activity, researchers may need to identify contexts in which the pupil-LC correspondence has been validated, and be aware of where it has not. In addition, there is a need to examine the potential contributions of other neuromodulatory systems that play critical roles in regulation of arousal, and their association with pupillary dynamics.

Critical challenges are therefore present in the attempt to use pupillary dynamics to understand the health of the LC, or any other brain region, in the context of aging. First, it remains necessary to validate experimental paradigms where pupil size reliably reflects LC activity. Second, aging-related changes may alter LC structure (Dahl et al., 2019; Liu et al., 2019; Riley, Cicero, Mabry, et al., 2025), activity (Hämmerer et al., 2017; Riley, Cicero, Swallow, et al., 2025), and connectivity (Cicero et al., 2025; Jacobs et al., 2015; Schneider et al., 2024), including with the brain regions that directly control pupil size. Last but not least, there are peripheral changes in pupillary control, e.g., age-related miosis and reduction of dynamic range (Larsson & Österlind, 1943; Riley et al., 2023; Winn et al., 1994), that may mask or complicate the relationship between aging, pupillary dynamics and the LC. The majority of studies characterizing the pupil-LC relationship have been conducted in younger populations, raising questions about whether findings can be generalized to aging cohorts. The relationship between pupillary and putative LC-NE system activity may be modified by age, beyond age-related peripheral changes.

To address these issues, we conducted a study combining functional MRI measurements of pupil derivative-BOLD coupling maps in 89 adults aged 19-78, while examining attentional orienting during an oddball task shown previously to elicit pupil-LC coupling (Murphy et al., 2014). We sought to evaluate the extent to which pupillary dynamics serve as a proxy for activity in multiple candidate neurochemical systems, derived from PET receptor and transporter density maps (Hansen et al., 2022; Markello et al., 2022), and whether these associations differ across the adult lifespan.

## Results

### 1. Task performance

Participants were asked to push a button when infrequent visual oddballs (20% of task stimuli) appeared on the screen. The response window was 1.5s. One participant responded to roughly 90% of all trials, including standard stimuli, and was thus excluded from the behavioral analysis. The remaining participants detected a mean of 86.0% (SD = 10.5%) of oddballs with a mean reaction time of 531 ms (SD = 63). False alarms to non-oddball (standard and bright) stimuli were rare (mean = 1.4%, SD = 1.1%). To test for age group differences, we fit a linear model on response accuracy with a fixed effect of age group. Oddball detection rate did not differ across age groups (younger 88.6%, middle 81.3%, older 88.3%; F[2,40] = 2.46, p = 0.10), but there was a marginal increase in the false-alarm rate in older adults (younger 1.1%, middle 1.5%, older 2.3%; F[2,40] = 2.99, p = 0.06). Reaction time did differ with age (F[2,40] = 3.43, p = 0.042, ηp² = 0.15), being slowest in older (M = 578 ms, SD = 59) and fastest in younger adults as expected (M = 511 ms, SD = 51; middle M = 538 ms, SD = 71).

### 2. Pupillary responses by trial type and age

We measured task-evoked pupillary responses to standard, oddball, and bright trials, with the latter serving as a task irrelevant low-level stimulus control condition. Responses were expressed as a percentage of each participant’s dynamic range, because dynamic range decreased significantly with age (younger M=2181 arbitrary units, middle M=1501, older M=1143; t(84) = −4.03, b=−29.0/year, p<.001), consistent with age related miosis. To evaluate how these task-evoked pupillary responses varied with stimulus type and age, we fit a linear mixed-effects model on peak pupillary dilation (as a percentage of dynamic range) with fixed effects of trial type and age group and their interaction, and a random intercept for participant. A parallel model was constructed for the area under the curve of pupillary responses (the integrated response, rather than simply the peak). Oddball events provoked robust pupillary dilations that were significantly larger than those to standard events (oddball peak = 23.5% of dynamic range, standard = 14.8%, bright = 13.3%; main effect of trial type F[2,21737] = 434.03, p < 0.0001; oddball > standard by 8.7 percentage points, t(21738) = 27.49, p < 0.0001; oddball > bright by 10.2 percentage points, t(21719) = 25.69, p < 0.0001, standard > bright by 1.5 percentage points, t(21737) = 4.85, p < 0.0001, **Fig. 1A**). Peak dilation also varied across age groups overall (F[2,85.1] = 3.33, p = 0.040), and, more importantly, trial type interacted with age (F[4,21737] = 8.47, p < 0.0001). This interaction was non-monotonic: the oddball-evoked dilation was largest in middle-aged (26.7%), intermediate in older (24.0%), and smallest in younger adults (19.7%), whereas standard and bright responses were comparable across groups. Within oddball trials, middle-aged adults dilated significantly more than younger adults (t(104) = 3.96, p < 0.001), whereas older adults did not differ reliably from either younger (t(105) = 2.16, p = 0.096) or middle-aged adults (t(105) = 1.35, p = 0.45); the interaction thus reflects a peak in middle age. The area under the curve showed the same pattern: a robust effect of trial type (F[2,7687] = 134.82, p < 0.0001; oddball > standard t(7687) = 15.48, oddball > bright t(7670) = 14.62, standard > bright t(7691) = 2.28, p = 0.058) and a trial-type-by-age interaction (F[4,7682] = 5.85, p = 0.0001), but no significant overall age effect (F[2,89.0] = 1.88, p = 0.16). As with peak dilation, the integrated oddball response was largest in middle-aged and smallest in younger adults (middle > younger, t(422) = 3.97, p < 0.001), with older adults intermediate and not significantly different from either other group (older vs. younger t(456) = 1.91, p = 0.16; older vs. middle t(383) = 1.61, p = 0.29).

**Fig. 1.**
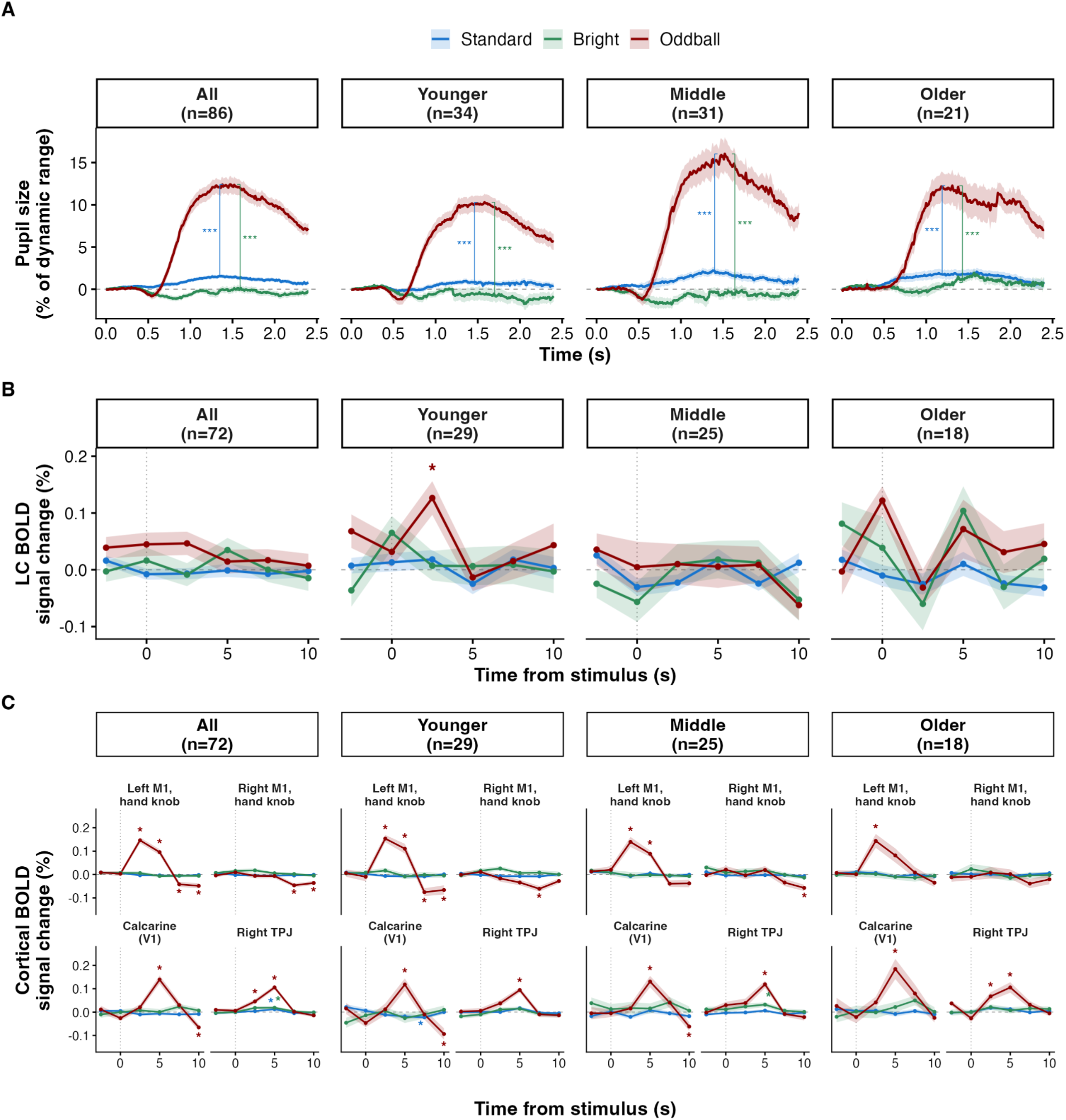
Task-evoked pupillary and locus coeruleus responses by age group. (A) Task-evoked pupillary response to each trial type (Standard, Bright, Oddball), shown for the whole sample (“All”) and by age group. Pupil size is expressed as a percentage of each participant’s dynamic range. (B) Locus coeruleus (LC) BOLD response to the same trial types, estimated with a finite impulse response (TENT) model at 2.5 s intervals from −2.5 to 10 s after stimulus onset within each participant’s SegmentAAN Bayesian LC label in native space, after regression of cardiac, respiratory, and 4th ventricle nuisance signals (n = 72; see Methods). The pre-stimulus (−2.5 s) and onset (0 s) estimates are shown as checks and were not treated as responses. (C ) Cortical BOLD responses to the same trial types, estimated using the same model as in B, at the hand knob of the left and right motor cortices, the calcarine cortex, and right temporoparietal junction, as positive controls (except hand knob of right motor cortex, which is a negative control). All lines show group means ± SE. Per-panel sample sizes are indicated. Asterisks mark significant results. For (A) they mark significant peak contrasts from the model in Section 2. For (B) and (C) they mark significant knots after correction from the model in Section 3.

### 3. Estimated hemodynamic responses in the LC

We estimated BOLD responses to standard, bright, and oddball events within each participant’s LC, expressed in % signal change (Fig. 1B). Because the LC occupies an average volume of just 5 voxels at 2 mm isotropic resolution, we combined voxels by partial volume-weighted averaging (i.e., weighting each voxel by its fractional overlap with each individual’s LC ROI, which was defined at 1 mm isotropic resolution). Responses were estimated with a finite impulse response (TENT) model that makes no assumption about response shape, on multi-echo denoised data with cardiac, respiratory and 4th ventricle nuisance regressors (n = 72). Within each age group, every condition’s response at each latency from 2.5 to 10 s was tested against zero with a single permutation-based maximum-statistic correction across the three conditions and four latencies. In younger adults (n = 29), the LC responded to oddballs at 2.5 s after stimulus onset (M = 0.126%, t(28) = 4.19, corrected p = .003). This was the only response to survive correction in any group. Younger adults showed no response to standard or bright stimuli, and middle-aged (n = 25) and older adults (n = 18) showed no significant response to any stimulus type (all corrected p > 0.27, despite averages far from zero at some knots in older adults). There were more younger adults than middle-aged or older adults in the sample; the younger adults’ effect size at the significant 2.5 s knot (d = 0.78, comparing the size of the observed peak with 0) gives 96% and 87% power to detect a comparable BOLD response at the middle-aged and older adult sample sizes, respectively.

The effect of each nuisance model on the response is shown in **Supplementary Fig. S2,** and the effect of the same nuisance regressors on modeled (SPMG3, as opposed to model-free FIR) hemodynamic responses is shown in **Supplementary Fig. S3**. The early peak in the LC is consistent with the faster, narrower hemodynamic response of subcortical structures to brief stimuli (Lewis et al., 2018). The 4th ventricle showed no oddball response in any group at any knot (all corrected p > 0.12).

To test the spatial specificity of the observed LC oddball-evoked response, we also examined oddball-evoked responses in other ascending arousal system nuclei (Olchanyi et al., 2024) with centroids within 10 mm of the LC (parabrachial complex, laterodorsal tegmental nucleus, oral pontine reticular formation, median and dorsal raphe), and in two control regions (a 1-4 mm shell of pontine tegmentum surrounding the LC and not sharing any full or partial voxels with the LC or 4th ventricle, and the 4th ventricle itself), using the same nuisance regressors. We found that the LC had the largest oddball-evoked response, but significant responses were also observed within the parabrachial complex, oral pontine reticular formation, and the shell of pontine tegmentum in younger adults only (**Supplementary Fig. S4**). The former two findings are not unexpected given the role of these regions in salience detection and orienting. The latter finding could be caused by registration challenges, hemodynamic spread, or simply true oddball-evoked activity in this area.

As controls, the same model was applied to Schaefer-200 representing the hand knobs of left and right primary motor cortex (±37, −25, 58; (Yousry et al., 1997)), calcarine (V1) cortex (±8, −85, 5; left and right parcels averaged) and right temporoparietal junction (54, −50, 20; (Krall et al., 2015)). Robust oddball-specific activity was observed in the hand knob of left, but not right, primary motor cortex, associated with button-pushing with the right hand. Robust oddball-specific activity was also observed in the right temporoparietal junction and V1, presumably associated with attentional orienting toward and visual processing of oddballs, respectively.

### 4. Whole-brain univariate analysis

To characterize whole-brain task activation, and test for effects of age group, we fit a voxelwise linear mixed-effects model on single-participant canonical HRF response amplitudes with fixed effects of trial type, age group and their interaction, and a random intercept for participant. For cluster thresholds, we used a voxelwise threshold of p < 0.001 and a family-wise error rate of p < 0.05, estimated by Monte Carlo simulation from the spatial autocorrelation of the single-participant residuals.

Trial type modulated the BOLD response across widespread cortical and cerebellar areas (main effect of trial type: peak chi-square = 59.5 at the left supramarginal gyrus; 6,937 voxels surviving correction, see **Supplementary Table S1**). Oddballs evoked greater activation than standards throughout canonical target-detection areas (bilateral supramarginal and inferior parietal cortex (left peak Z = 6.8 at MNI [−63, −43, 39]; right peak Z = 6.6 at [59, −37, 45]), lateral and dorsomedial frontal cortex, and cerebellum (5,871 voxels), together with deactivation of midline posterior cingulate and precuneus (oddballs < standards; e.g., Z = −5.8 at [−5, −43, 37]). Bright stimuli did not differ from standards anywhere after cluster correction.

Oddball-evoked activation also varied with age. Older adults showed larger oddball responses than younger adults across midline sensorimotor and default mode regions: paracentral lobule, anterior cingulate, precuneus, and thalamus (younger < older; 587 voxels in 55 clusters; peak Z = −5.6 at the right precuneus). However, the standard response did not differ by age group, the bright response differed only in four small clusters (34 voxels, all younger < older; **Supplementary Table S1**), there was no main effect of age group, and the trial-type-by-age group interaction resulted in only small clusters (17 voxels total) after cluster correction. In summary, robust oddball activation was accompanied by comparatively weak effects of age group, which were largely specific to the oddball condition.

### 5. Whole-brain pupil-BOLD coupling (amplitude-modulated GLM)

To test whether the trial-by-trial BOLD response covaried with pupillary responses to those trials, we fit single-participant GLMs with amplitude-modulated trial type regressors and then carried out the same voxelwise mixed-effects model and cluster correction used in Section 4 (see **Supplementary Table S2**). Three pupil modulators were used: pretrial pupil size, area under the curve of individual task-evoked pupillary response for each trial, and the peak rate of change of pupil diameter during each trial (in % of dynamic range per second, i.e., maximum derivative of pupil size). In all three analyses, very few voxels appeared to have significant pupil-BOLD coupling. The pupil size model (i.e, with pretrial pupil size as the modulator) yielded 736 voxels distributed across 80 clusters (largest 29 voxels). The area under the curve model yielded 85 voxels across 10 clusters (largest 13 voxels), and the derivative model yielded 98 voxels across 13 clusters (largest 11 voxels). The largest clusters fell in occipital and calcarine cortex, with the remaining clusters scattered across frontal, parietal, temporal, and cerebellar cortex. None of the three modulators produced a cluster overlapping with the LC (either the Bayesian or hand-traced ROI).

### 6. Pupil-bold coupling with a continuous task-naive regressor

To test whether our data reproduce widespread pupil-BOLD coupling (Murphy et al., 2014), we replaced the trial-wise amplitude-modulated model with a continuous-regressor approach: each individual’s continuous pupil time series and its first derivative were downsampled to the TR rate (one value per TR), convolved with a canonical HRF, and entered directly as regressors in a GLM. We then constructed voxelwise linear mixed-effects models on the single-participant pupil-regressor coefficients with fixed effects of pupil regressor type (either pupil size or its derivative) and age group and their interaction, and a random intercept for participant. This approach was task-naive and did not account for trial conditions. See **Supplementary Table S3** for full results. The temporal derivative of pupil size produced a large, predominantly positive coupling field (**Fig. 2B**): 24,948 voxels in 132 clusters, of which 103 (24,489 voxels) were positive, dominated by one contiguous 22,087-voxel positive cluster (peak Z = 9.26, left lingual gyrus) spanning occipital, parietal, insular, striatal, cerebellar, and brainstem areas. Pupil size (**Fig. 2A**) itself yielded a small, predominantly negative field of 214 voxels in 18 clusters (17 negative; largest 44 voxels, right precentral gyrus), distributed across occipital, temporal, and sensorimotor cortex. The pupil derivative result was broadly consistent with Murphy et al., 2014; the pupil size result also matches previous reports (DiNuzzo et al., 2019). This coupling showed little evidence of age dependence. The main effect of age group produced no surviving clusters, and the regressor-by-age group interaction produced only two small clusters (12 and 7 voxels). Pupil-size and pupil-derivative coupling with BOLD therefore did not appear to vary meaningfully across age groups in this analysis.

**Fig. 2.**
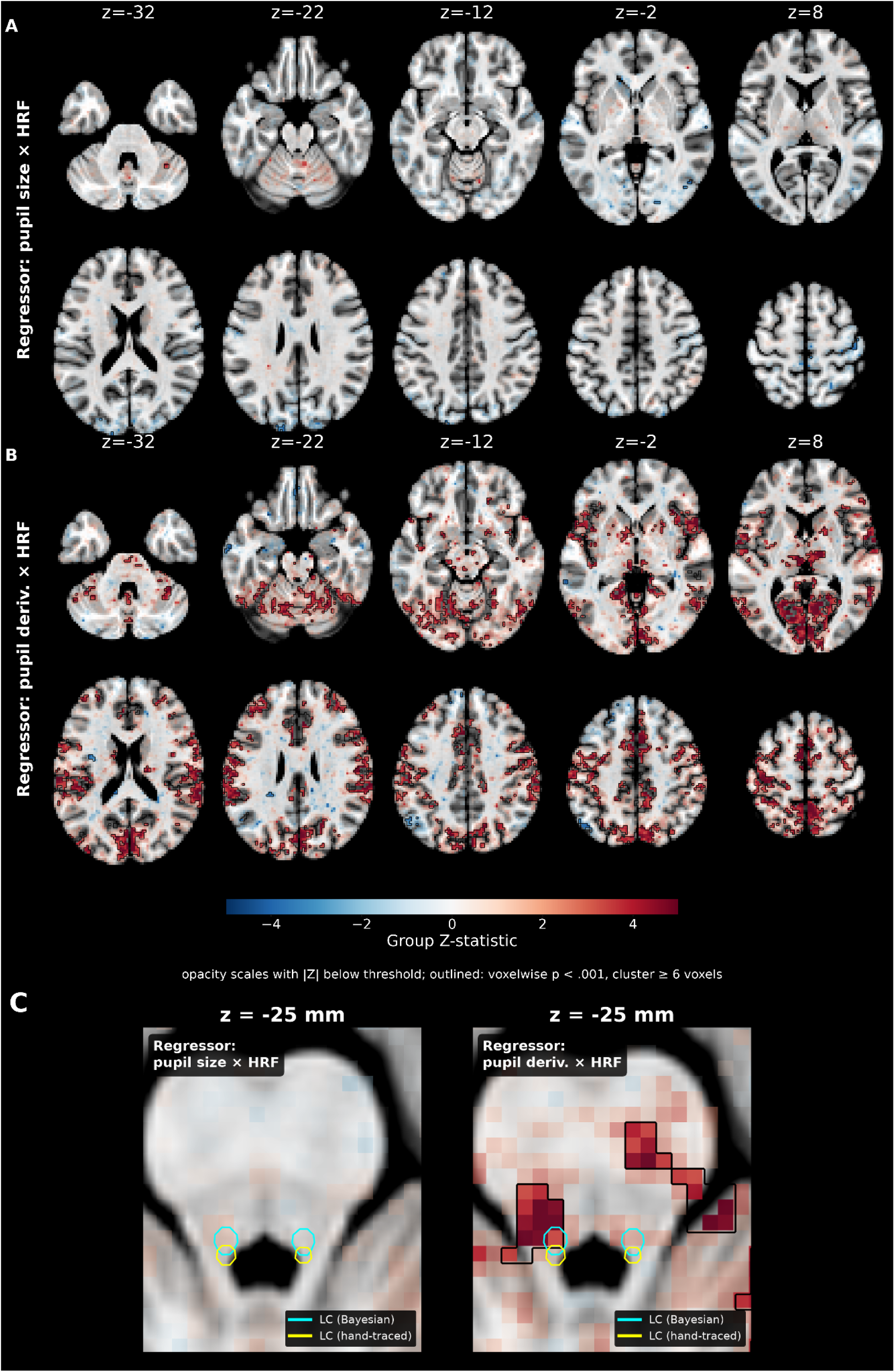
Whole-brain pupil-BOLD coupling. Voxels whose BOLD signal significantly covaries with the pupil size (A) and pupil derivative (B) regressors (each convolved with the hemodynamic response), N = 81. (C) Brainstem detail at two axial levels, with Bayesian (cyan) and hand-traced (yellow) LC ROIs. Significant voxels (black outlines) at p < 0.001, cluster ≥ 6 voxels.

To test whether either set of results (i.e., either significant pupil size or pupil derivative coupling with BOLD) intersected with the LC, we tested the overlap between significant pupil-brain voxels and our two LC ROIs (Bayesian and hand-traced). Pupil size coupling did not significantly overlap with the LC; pupil derivative coupling overlapped by at most two voxels (Bayesian ROI: 2 of 11 voxels; hand-traced ROI: 0 of 8, **Fig. 2C**). However, a significant cluster of pupil derivative-BOLD coupling lay very close to the LC: there was a strong brainstem cluster (peak Z = 6.18) 2-4 mm away from the LC centroid in the dorsal pons. This distance is within the potential error of nonlinear alignment of brainstem regions from individual space to MNI space. In summary, this continuous-regressor analysis reproduces widespread pupil-derivative-BOLD coupling, including robust brainstem coupling in the immediate vicinity of the LC.

### 7. Pupil-BOLD cross-correlation

In previous analyses, we identified significant task-evoked pupillary responses, task-evoked hemodynamic responses in the LC in younger adults, and significant pupil-BOLD coupling. The latter included strong brainstem clusters adjacent to our LC ROIs. However, the LC response was detectable only at the group level; individual LC response estimates were too variable to serve as a per-person measure. Therefore, we next attempted to characterize areas of the cortex that covary with pupil size or derivative. First, because we were uncertain about the precise length of the time delay between pupillary responses and BOLD responses, we computed the cross-correlation between the continuous pupil and pupil derivative time series (each separately) and the mean BOLD of each Yeo-7 resting-state network across pupil-to-BOLD delays of −2 to +10 s (step size 0.25 s, **Fig. 3**). Across the entire sample, the correlation coefficient peaked at +2.25 s for pupil size (pupil size 2.25 s ahead of BOLD) and +5.00 s for pupil derivative (change in pupil size 5.00 s ahead of BOLD). The peak of the pupil derivative-BOLD cross-correlation did not differ significantly between age groups (4.8, 5.4, and 5.7 s for younger, middle-aged, and older adults; F[2,78] = 1.73, p = 0.18, **Fig. 3I**). The pupil size-BOLD peak occurred later in older adults (1.6, 2.1, and 2.7 s for younger, middle-aged, and older adults, F[2,78] = 4.88, p = 0.010). However, for both pupil-BOLD and pupil derivative-BOLD cross correlation, individual correlation strengths at the group average delay, expressed as a fraction of each participant’s own maximum, did not differ across age groups (pupil size: 82%, 89%, and 90% of maximum, p = 0.60; pupil derivative: 72%, 68%, and 72% retained, p = 0.80). Therefore, going forward, when measuring pupil-brain coupling, we used these fixed time delays (2.25 and 5.0 s).

**Fig. 3.**
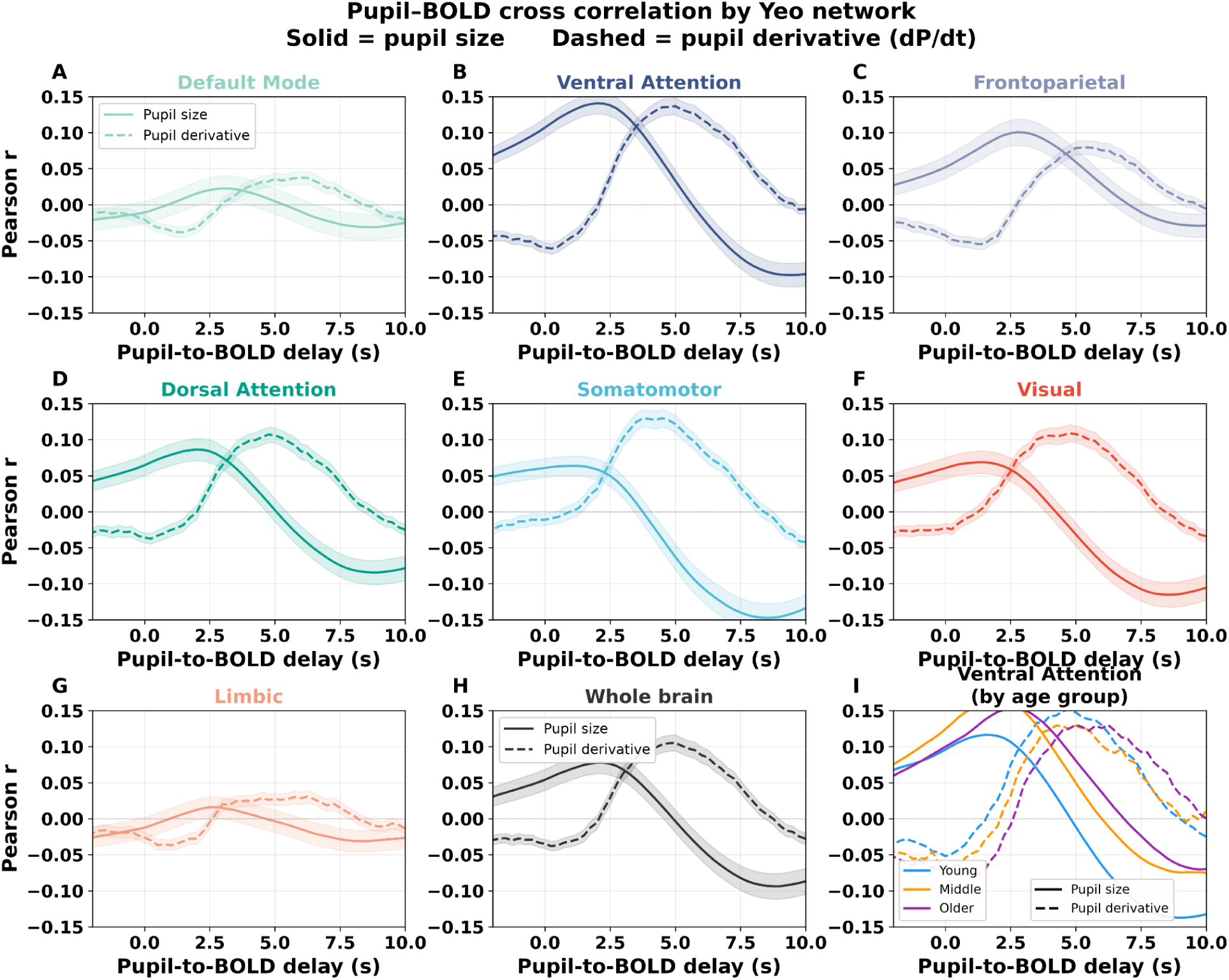
Pupil-BOLD cross-correlation across time delays between −2 to +10 s. Cross-correlation (r) between pupil and mean BOLD signal as a function of time delay, for each of the seven Yeo networks, the whole brain, and the ventral attention network (highest peak r in full sample) split by age group. Solid lines = pupil size; dashed = pupil derivative. Shaded bands show ± SE (N = 81).

To determine how pupil-BOLD coupling changed by cortical region and by age group, we fit linear mixed-effects models on the pupil-BOLD and pupil derivative-BOLD correlation coefficient within each voxel with fixed effects of Yeo network and age group and their interaction, and a random intercept for participant. Coupling between both pupil size and pupil derivative and BOLD varied significantly across Yeo networks (pupil size: F[6,468] = 17.70, p < 0.0001; pupil derivative: F[6,468] = 55.05, p < 0.0001). The Ventral Attention network had the highest correlation for both pupil size (r = 0.142) and pupil derivative (r = 0.136), followed by Frontoparietal for pupil size (r = 0.103) and Somatomotor for pupil derivative (r = 0.118); Limbic and Default Mode networks were weakest for both (r ≤ 0.036). Neither the model for pupil size-BOLD nor pupil derivative-BOLD coupling had a significant main effect of age group (size: F[2,78] = 2.84, p = 0.064; derivative: F[2,78] = 0.04, p = 0.96). However, the network-by-age group interaction was significant for pupil derivative (F[12,468] = 2.19, p = 0.011), but not for pupil size (F[12,468] = 0.78, p = 0.67). Following up on this interaction, we found that pupil derivative-BOLD coupling appeared to be weaker in older adults than younger adults in sensory/attention networks (Somatomotor Δ = −0.043, Visual Δ = −0.022, Ventral Attention Δ = −0.017, Dorsal Attention Δ = −0.013), was essentially unchanged in the Default Mode network (Δ = −0.001), and rose in the Limbic (Δ = +0.028) and Frontoparietal (Δ = +0.021) networks. However, no individual within-network age contrast survived Tukey correction (all pairwise p > 0.24).

For each participant, we computed a whole-brain coupling map for pupil size (+2.25 s) and pupil derivative (+5.00 s). Then, we tested each voxel against zero across all participants to determine which voxels were significantly correlated with pupil size or pupil derivative (**Fig. 4A,B**). Pupil size was significantly coupled to BOLD in 14,078 voxels in 49 clusters (38 positive, totalling 13,734 voxels), with the largest clusters in cerebellum (5,162 voxels, peak t = 7.96), middle cingulate and precuneus (peak t = 8.20 at the middle cingulate), and bilateral supramarginal cortex (right 1,191 voxels, t = 7.30; left 737 voxels). Pupil derivative was significantly coupled to BOLD in 33,629 voxels in 25 clusters (24 positive, totalling 33,611 voxels), dominated by a contiguous 31,812-voxel cluster (peak t = 8.85 at the left precuneus) that spanned parietal, cingulate, frontal, insular, striatal, cerebellar, and brainstem areas.

**Fig. 4.**
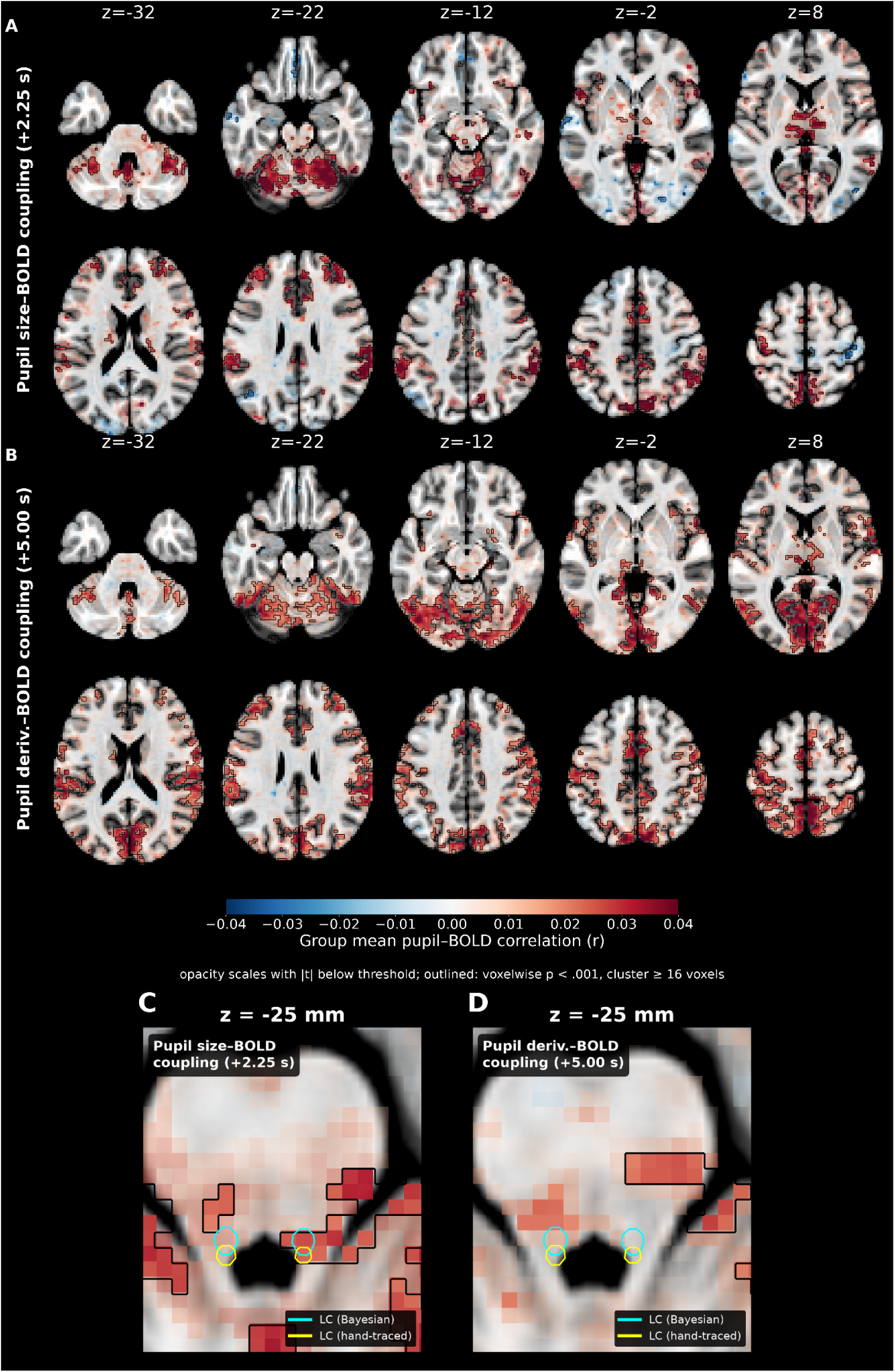
Pupil-BOLD coupling at time of peak correlation. Whole-brain coupling evaluated at the time of each regressor’s peak correlation within the whole sample: pupil size at +2.25 s (A) and pupil derivative at +5.00 s (B), with brainstem detail and Bayesian (cyan) and hand-traced (yellow) LC ROIs for pupil size (C ) and pupil derivative (D). Significant voxels (within black outlines) are derived from a group one-sample t test vs zero, p < 0.001, cluster ≥ 16 voxels (N = 81).

The pupil derivative-BOLD coupling map was strongly correlated with the continuous-regressor (task-naive) derivative coupling map from Section 6. Dividing both maps into Schaefer 200 parcels, we found a very strong correlation (parcelwise r = 0.95, spin p = 0.0002). The pupil size coupling map had a weaker, but still substantial, correlation with its counterpart from Section 6 (parcelwise r = 0.66, spin p = 0.0002). This is most likely because the Section 6 analysis used the same canonical HRF, peaking at 5s, for both pupil size and pupil derivative. Under those conditions, the unique contribution of pupil size in explaining BOLD activity is smaller and negative. However, the cross-correlation analyses described in this section resulted in different time delays being used for pupil size (2.25 s) vs pupil derivative (5 s), a schema under which both features show positive correlations.

The correlation between pupil derivative and BOLD was, by definition, associated with moments of rapid pupil size change. We tested whether these moments were associated with the oddball task, or whether they were spontaneous. We identified the 60 highest-derivative events (positive derivative only, after downsampling to 4 Hz to avoid accidental identification of noise) within each participant’s pupil timeseries, because there were 60 oddball events in the task. Then we compared the timing of these high-derivative events to the 2 s windows after each oddball event, asking whether high-derivative events were oddball events. Oddball events preceded highest-derivative events 1.4 times more often than expected under a null distribution (i.e., shifting the onset times of oddball events by a random offset and repeating 1000x; 21.1% of highest-derivative events occurred after true oddball events vs 14.9% in the null distribution; paired t(85)=3.63, p=0.0005). Standard trials showed no such effect (50.3% vs 51.1%). Bright trials had significantly fewer highest-derivative events in the true data than in the null distribution (11.2% vs. 15.2% expected; paired t(85)=-6.85, p < 0.001). This is consistent with pupil constrictions caused by the bright stimulus taking place instead. In total, we found that moments of rapid pupil size change were significantly, but not exclusively, associated with the oddball task.

We next tested whether the significant cross-correlation coupling maps intersected with our two LC ROIs **(Fig. 4C,D)**. Significant pupil size-BOLD coupling (+2.25 s) overlapped with the LC (Bayesian ROI: 3 of 11 voxels; hand-traced ROI: 1 of 8), whereas significant pupil derivative-BOLD coupling (+5.00 s) did not overlap either ROI (Bayesian: 0 of 11; hand-traced: 0 of 8); its nearest suprathreshold voxel was 6.3 mm (Bayesian) or 7.5 mm (hand-traced) from the LC. However, a significant cluster near the left LC, which did not pass the 16-voxel cluster threshold, was noted (10 voxels, 2.0 mm from the left LC; cluster-mean r = 0.021, t(80) = 5.73, p < .001). These overlap patterns were stable across LC ROI probability cutoffs from 0.05 to 0.35. The cross-correlation profile of the partial-volume-weighted LC ROI time series itself is shown in **Supplementary Fig. S5**. The cross correlation profiles within the LC resembled those observed in cortex, with an earlier peak of pupil size coupling and a later peak of pupil derivative coupling, both of which were significant.

To determine whether pupil-BOLD coupling differed between age groups, we quantified each participant’s coupling extent (count of voxels with r > 0.10 per participant, chosen because group size and thus threshold t-statistics differed between groups) and also each participant’s mean correlation (mean r across all voxels in the brain). For pupil derivative, BOLD coupling extent was higher in younger adults (median voxels with r > 0.10: 10,803, 7,978, and 5,514 voxels for young, middle-aged, and older, F[2,77] = 3.17, p = 0.047; young-versus-older t = 3.06, p = 0.004, linear age effect r = −0.22, p = 0.055, **Fig. 5B**). Threshold-free mean r followed the same young > middle-aged > older trend but was not significant (full F[2,78] = 0.45, p = 0.636, **Fig. 5D**). Pupil size-BOLD coupling showed no age effect on either metric (extent F[2,78] = 0.22, p = 0.80; mean r F[2,78] = 1.06, p = 0.35, **Fig. 5A&C**). The association between age and coupling extent differed significantly between pupil size vs. pupil change (r = +0.03 for pupil size versus r = −0.22 for pupil derivative; Williams’s test for dependent correlations, t[77] = 2.07, p = 0.041).The relatively mild effect of age thus appeared to be specific to pupil derivative-BOLD coupling, and to how much cortex is recruited (extent) more than per-voxel magnitude. Younger adults had more cortical voxels significantly correlated with pupil derivative than older adults.

**Fig. 5.**
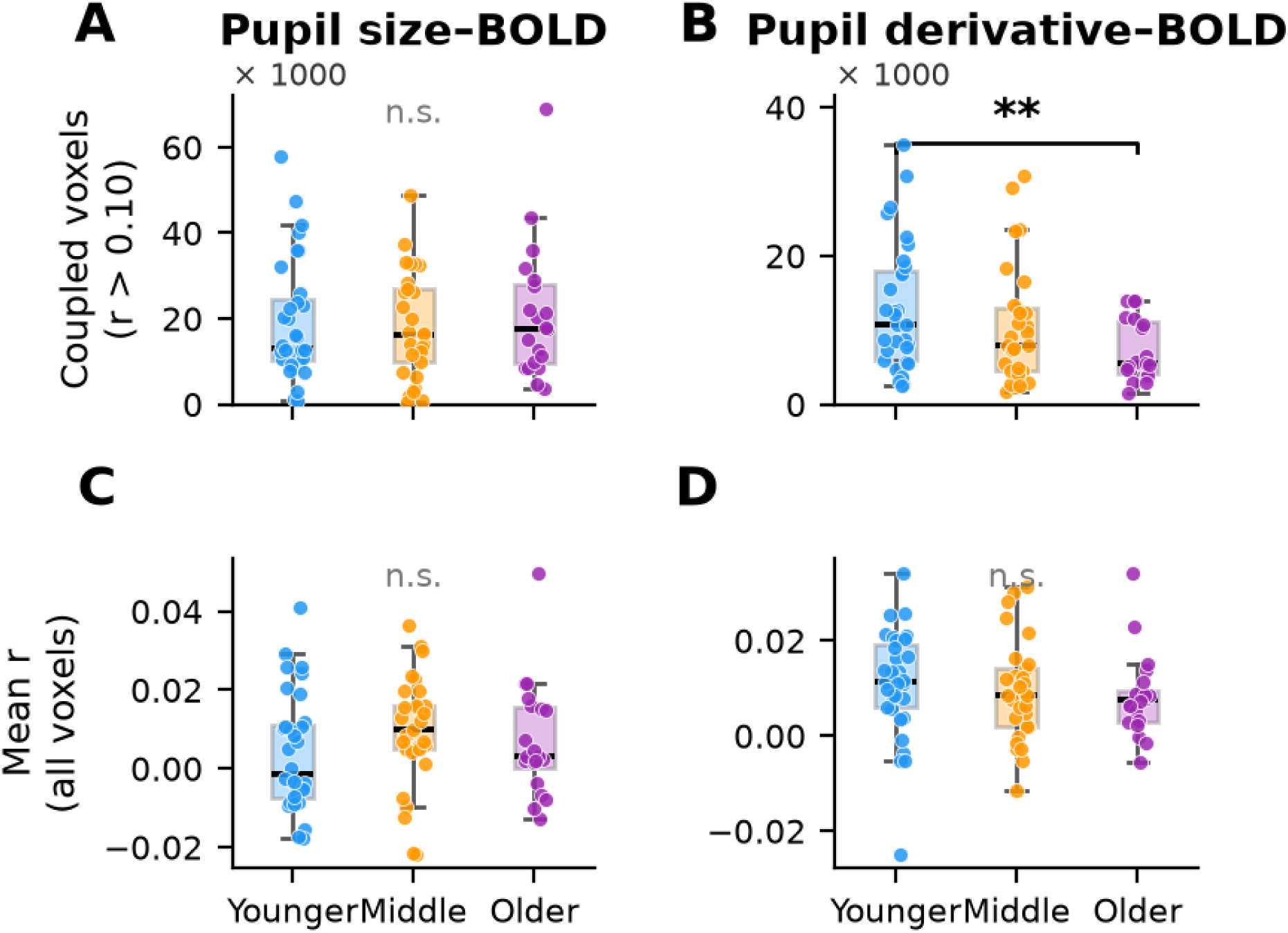
Pupil-BOLD coupling by age group, quantified as whole brain spatial extent (voxels with fixed-lag r > 0.10; A, B) and threshold-free mean r across all in-brain voxels (C, D), for pupil size (left) and pupil derivative (right). Boxes show medians and IQRs; points are individual participants. Pupil derivative-BOLD coupling extent decreases with age (** = Younger > Older, Welch p = .004).

To confirm that these coupling maps reflected a true pupil-related relationship, we independently cross-correlated heart rate, heart rate variability (HRV) and respiration with whole-brain BOLD (after multi-echo denoising) using parallel pipelines; heart rate and respiration rate did couple with BOLD. We thus recomputed voxelwise pupil-BOLD coupling maps after regressing out heart rate, respiration rate and HRV from both the BOLD and the pupil regressors. These analyses demonstrated that the spatial patterns of pupil-BOLD coupling were not attributable to cardiac or respiratory fluctuations (see **Supplement and Supplementary Fig. S1**).

Finally, because the pupil size-BOLD and pupil derivative-BOLD coupling maps overlapped spatially, we asked how much of the derivative-coupling pattern was independent of pupil size-coupling pattern. Parcellating both group t-maps into Schaefer-200 regions, the two maps were positively correlated in space (r = +0.607, spin-test p = 0.0002), sharing 36.9% of their spatial variance. We constructed residualized-derivative and residualized-size maps (isolating each component by removing the other) by regressing pupil size out of the pupil derivative parcels, and the pupil derivative parcels from pupil size (residual r = 0.000, by design) for use in further analysis.

### 8. Spatial association of pupil-BOLD coupling with neurotransmitter density

To determine whether the whole-brain pupil size-BOLD and pupil derivative-BOLD coupling maps were associated with particular neurotransmitter system, we employed 19 PET receptor/transporter density maps spanning 9 systems (acetylcholine, dopamine, endocannabinoid, GABA, glutamate, histamine, norepinephrine, opioid, and serotonin, all available publicly through neuromaps (Markello et al., 2022)), and measured their Schaefer-200 parcel-level spatial correlation with pupil size-BOLD and pupil derivative-BOLD coupling t-maps. Pupil size-BOLD coupling did not spatially correlate with any neurotransmitter density maps (all q > 0.30).

However, partialling pupil derivative-BOLD coupling out of pupil size-BOLD coupling (i.e., isolating the pupil size-unique coupling pattern) revealed significant spatial correlations with μ-opioid receptor density (MOR; r = +0.578, spin p < 0.001, FDR q = 0.004), cannabinoid CB1 density (r = +0.430, p = 0.002, q = 0.015) and metabotropic glutamate receptor 5 density (mGluR5; r = +0.376, p = 0.003, q = 0.018). No other PET maps were significantly correlated.

Pupil derivative-BOLD coupling was significantly correlated with norepinephrine-transporter (NET) density (**Fig. 6**). NET had the highest positive spatial correlation with pupil derivative-BOLD coupling and was the only map surviving Benjamini-Hochberg FDR correction (r = +0.422, spin p = 0.001, q = 0.027). The next-ranked tracers were two serotonergic maps reaching only uncorrected significance (5-HT4, 5-HT1a, both negative correlations; FDR q > 0.19). NET density was also significantly correlated with the continuous-regressor (task-naive) GLM derivative Z-map (r = +0.421, spin p < 0.001) from Section 6. Partialling the pupil size-BOLD coupling effect out of the pupil derivative-BOLD coupling parcels (i.e., isolating the relationship between pupil derivative and BOLD) strengthened the NET spatial correlation ( r = +0.502, spin p = 0.004). It also introduced new significant *negative* correlations with μ-opioid (MOR; r = −0.626, spin p = 0.001, q = 0.023) and cannabinoid (CB1; r = −0.555, spin p = 0.005, q = 0.030) receptor density, and strengthened one of the serotonergic negative correlations into FDR significance (5-HT1a r = −0.554, spin p = 0.006, q = 0.030). This is an expected result of the residualization process.

**Fig. 6.**
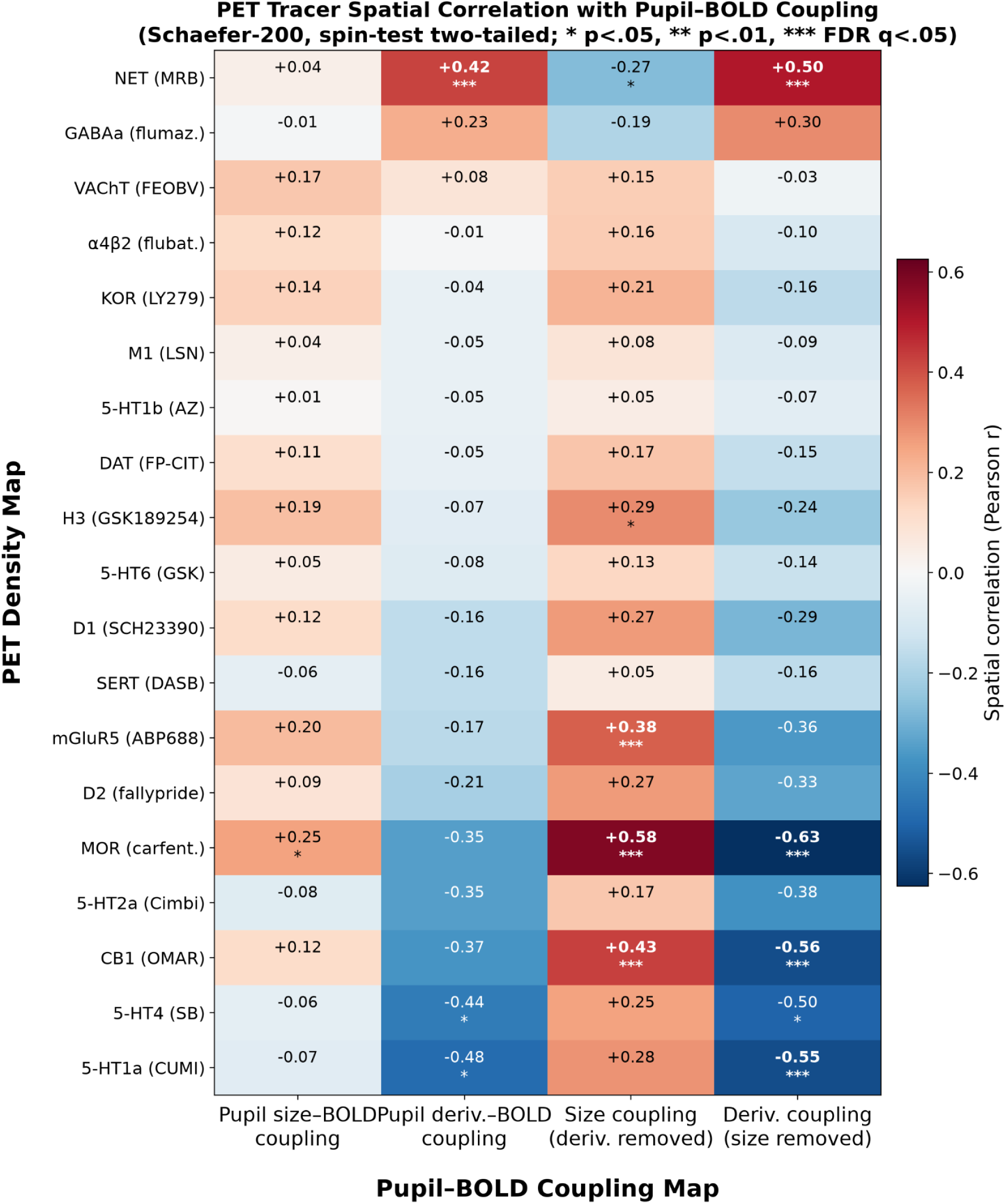
Relationships between pupil-BOLD coupling and neurochemical systems. Spatial correlation (Pearson r) between pupil-BOLD coupling maps (size, derivative, and each residualized for the other) and 19 neurotransmitter receptor/transporter PET density maps within Schaefer-200 parcels. Null distributions are obtained using an Alexander-Bloch spin test with 5,000 permutations. * p < 0.05, ** p < 0.01, *** FDR q < 0.05. Neurotransmitter density maps were drawn from Hansen et al., 2022, Jaworska et al., 2020, and Vijay et al., 2018 via neuromaps (Markello et al., 2022). Maps were, in the format of (target; tracer): the norepinephrine transporter (NET; [¹¹C]MRB); the dopamine transporter (DAT; [¹²³I]FP-CIT) and the dopamine D₁ (D1; [¹¹C]SCH23390) and D₂/D₃ (D2; [¹⁸F]fallypride) receptors; the serotonin 5-HT₁ₐ (5-HT1a; [¹¹C]CUMI-101), 5-HT₁ᵦ (5-HT1b; [¹¹C]AZ10419369), 5-HT₂ₐ (5-HT2a; [¹¹C]Cimbi-36), 5-HT₄ (5-HT4; [¹¹C]SB207145) and 5-HT₆ (5-HT6; [¹¹C]GSK215083) receptors and the serotonin transporter (SERT; [¹¹C]DASB); the vesicular acetylcholine transporter (VAChT; [¹⁸F]FEOBV) and the muscarinic M₁ (M1; [¹¹C]LSN3172176) and nicotinic α₄β₂ (α4β2; [¹⁸F]flubatine) acetylcholine receptors; the metabotropic glutamate receptor 5 (mGluR5; [¹¹C]ABP688); the GABA_A receptor (GABAa; [¹¹C]flumazenil); the cannabinoid CB₁ receptor (CB1; [¹¹C]OMAR); the histamine H₃ receptor (H3; [¹¹C]GSK189254); and the μ-opioid (MOR; [¹¹C]carfentanil) and κ-opioid (KOR; [¹¹C]LY2795050) receptors.

Next, we tested whether the spatial correlations between pupil-BOLD coupling and neurotransmitter receptor and transporter density were present within each age group, and whether they differed between age groups (**Fig. 7**). *Pupil size*. Significant spatial correlation between pupil size-unique BOLD coupling and MOR density was present in each age group (group-level parcelwise r = +0.554 younger, +0.461 middle, +0.528 older; all spin p = 0.0002). At the individual level the strength of the correlation did not differ significantly between age groups (ANOVA on Fisher-z F[2,78] = 2.33, p = 0.10; younger-versus-older Welch t = 0.49, p = 0.63; linear age effect r = −0.07, p = 0.54). The same was true for the spatial correlation with CB1 density. There was a significant positive correlation within each age group (group-level parcelwise r = +0.400 younger, +0.363 middle, +0.377 older; all spin p ≤ 0.009), and at the individual level (per-participant mean r = +0.14 younger, +0.07 middle, +0.09 older) there was no difference between age groups (ANOVA F[2,78] = 1.38, p = 0.26; linear age r = −0.09, p = 0.40). The spatial correlation with mGluR5 density was weaker and showed a non-significant trend toward age dependence. The group-level correlation was significant in younger and middle-aged adults (parcelwise r = +0.423, spin p = 0.003; r = +0.290, spin p = 0.010) but not in older adults (r = +0.253, spin p = 0.089). At the individual level (per-participant mean r = +0.13 younger, +0.05 middle, +0.06 older) age groups did not differ significantly (ANOVA F[2,78] = 2.72, p = 0.072; younger-versus-older Welch t = 1.56, p = 0.13; linear age r = −0.18, p = 0.12).

**Fig. 7.**
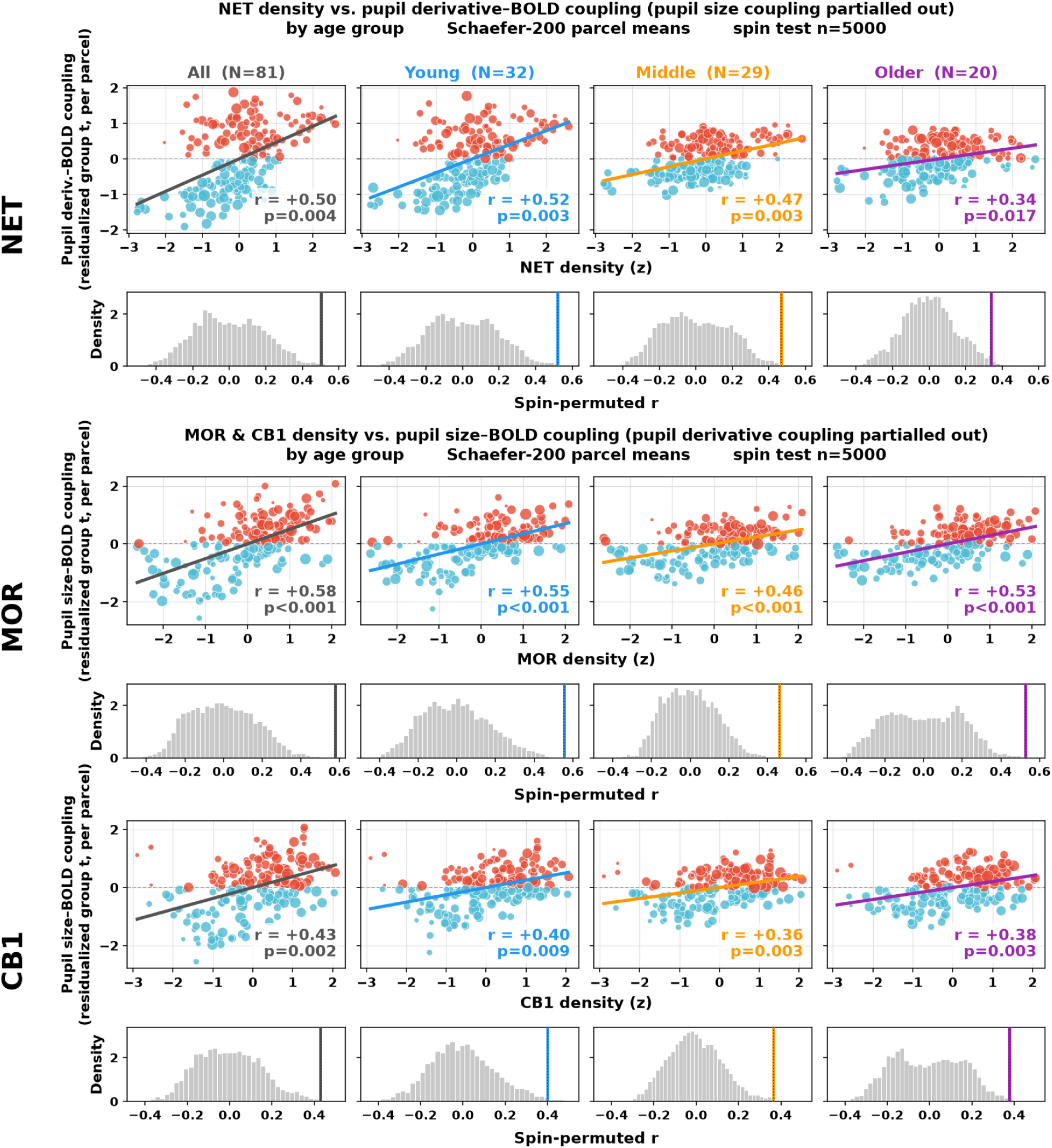
Pupil-BOLD coupling vs. significant neurochemical associations by age group. Residualized pupil derivative-BOLD correlation with PET NET density (top), and residualized pupil size-BOLD correlation with PET MOR and CB1 density (middle and bottom), shown for the whole sample and by age group. mGluR5 not shown. Each scatter is paired with its Alexander-Bloch spin-permutation null distribution (n = 5000).

*Pupil derivative*. Pupil derivative-BOLD coupling correlation with NET density differed between age groups. Significant spatial correlation between pupil derivative-BOLD coupling and NET density was still present in every age group (group-level parcelwise r = +0.519 younger, +0.468 middle, +0.338 older; spin p = 0.003, 0.003, and 0.017 respectively), but was significantly weaker in older than younger adults (per-participant mean r = +0.21 younger, +0.16 middle, +0.11 older, ANOVA on Fisher-z F[2,78] = 3.15, p = 0.048; younger-vs-older Welch t = 2.51, p = 0.016; linear age effect r = −0.28, p = 0.012). Despite this, within each age group, most participants had a positive (> 0) spatial correlation between pupil derivative-BOLD coupling and NET density: 87.5% of young adults, 89.7% of middle aged, and 80% of older adults.

Having found that NET density alone was positively correlated with pupil derivative-BOLD coupling, we asked whether the spatial organization shared by the two maps corresponds to a known axis of cortical organization. The principal gradient of resting-state functional connectivity (Margulies et al., 2016) is the dominant axis of this type, represented at one end by unimodal sensorimotor cortex and at the other by transmodal cortex. We first computed the spatial correlation between each of the 19 PET maps and this gradient. NET was the most strongly anticorrelated of the 19 (r = −0.49; the next-ranked, VAChT, was −0.17), meaning that NET density is highest in unimodal cortex and lowest in transmodal cortex. We then tested whether pupil derivative-BOLD coupling shared this organization. Both pupil derivative-BOLD coupling, and the version of the map with pupil size-BOLD coupling partialled out, were strongly anticorrelated with the principal gradient (r = −0.725, p = 0.0002, and r = −0.835, p = 0.0002 respectively). This was not the case for pupil size-BOLD coupling (r = −0.1, p = 0.40), dissociating the two metrics. Given this, we tested whether the NET correlation survived partialling the principal gradient out of both maps. It did not. The principal gradient accounts for more than half the spatial variance in derivative-BOLD coupling, so removing it removes a feature shared by NET and the coupling map. Notably, the MOR correlation with pupil size-unique coupling did survive the same procedure (r = 0.40, p = 0.0002 after gradient removal), as did the CB1 association (r = 0.25, p = 0.014 after gradient removal), indicating that this analysis is capable of detecting neurochemical alignment independent of the principal gradient.

Together these results suggest that areas of the brain in which BOLD covaries with change in pupil size (but not pupil size itself) are those richest in NET and most anticorrelated with the principal gradient. Areas in which BOLD uniquely covaries with pupil size are those with the most μ-opioid receptors, and secondarily cannabinoid CB1 and glutamatergic mGluR5 receptors. The extent of the correlation between change in pupil size and BOLD activity was significantly lower in older adults, as was the strength of the correlation between pupil change-BOLD coupling and NET density, although both remained significant. By contrast, neither pupil size-unique BOLD coupling nor the strength of spatial correlation between pupil size-BOLD coupling and opioid, cannabinoid, or mGluR5 receptors differed between age groups.

### 9. Individual differences in pupil reactivity

Last, we investigated whether individual differences in pupillary responses predicted individual differences in LC BOLD, pupil-brain coupling, or pupil-brain coupling’s association with NET, MOR, or CB1 density. For each participant, we used the peak oddball pupillary response as our pupil metric. Because pupillary responses vary non-monotonically with age (peaking in middle age), we adjusted for age group rather than treating age as a linear covariate. Oddball pupillary responses were centered within each age group and a single correlation was computed across the pooled centered values, equivalent to a partial correlation controlling for age group. We found no significant relationship between pupil oddball responses and oddball LC BOLD responses at 2.5 s (within-age r = +0.12, p = 0.32, n = 72, **Fig. 8A**). Because the LC response is a group-level result whose per-person estimate is dominated by estimation noise in a small ROI, individual-difference questions may be better addressed using whole-brain coupling metrics. However, we found that people with larger oddball-evoked pupil responses had pupil derivative-BOLD coupling more strongly correlated with NET density (within-age r = +0.24, p = 0.032, **Fig. 8B**), an effect that was strongest in younger adults but consistent in direction across all age groups. This association was relatively specific; oddball-evoked pupil responses did not significantly predict the correlation between pupil size-unique BOLD coupling and MOR (r = −0.06, p = 0.57) or CB1 (r = −0.03, p = 0.78) density, the two receptor systems with which pupil size-BOLD coupling was associated (Section 8). The more a person’s pupil reacted to oddballs, the more the change in their pupil size was associated with BOLD activity in NET-rich regions.

**Fig. 8.**
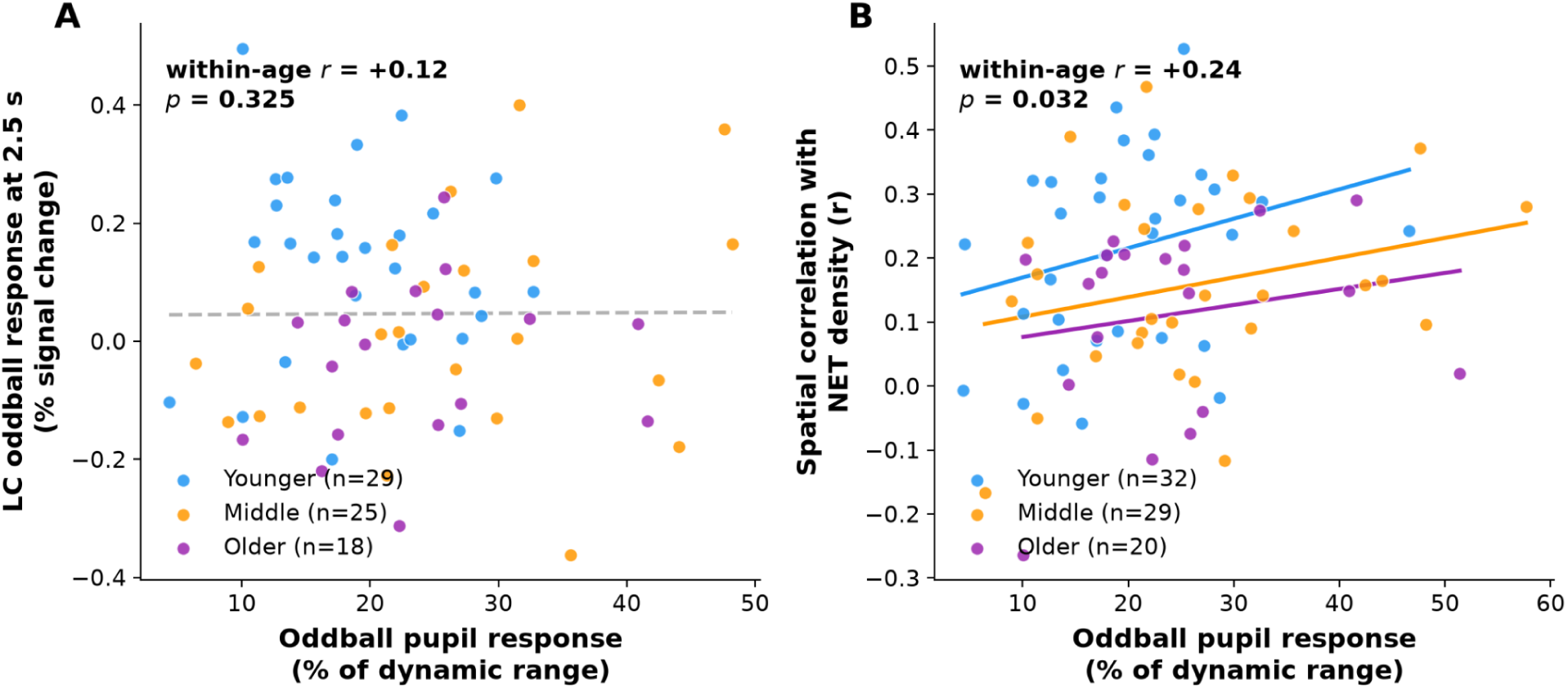
(A) Relationship between each participant’s oddball pupil peak (% of dynamic range) and LC BOLD oddball response at 2.5 s (finite impulse response estimate within individual LC ROIs in native space, nuisance-corrected), and (B) relationship between each individual participant’s oddball pupil peak and spatial alignment of pupil derivative-BOLD coupling with NET density, with within-age-group regression lines.

## Discussion

We observed robust task-evoked pupillary responses to oddball events in all age groups. Replicating our prior work (Riley et al., 2023) in a new cohort and task, we found that pupillary responses were largest in middle-aged adults compared to both younger and older adults. Within the LC, only younger adults showed a condition-specific oddball response, peaking 2.5 s after the stimulus. This replicates the age pattern of our previous work, in which orienting responses in the LC were detected in younger adults but not middle aged or older adults (Riley, Cicero, Swallow, et al., 2025), using a different task with three times as many target trials (192 vs 60). The absence of a detectable response in older adults, despite intact pupillary responses, is not explained by data quality, which was highest in the LC of older adults, although atrophy-related partial-volume dilution of the LC label cannot be ruled out as an explanation for this effect. Because the LC response is measurable at the group level but not reliably per person, we proceeded to whole-brain analyses of the relationship between pupil and BOLD activity. Using an approach inspired by Murphy et al., 2014, we created separate task-naive regressors for both pupil size and change in pupil size. We found that pupil change and pupil size were positively associated with activity in clusters in the immediate vicinity of the LC, at their respective delays. Pupil change was also associated with activity spanning the occipital and parietal cortex, insula, striatum, and cerebellum. These results broadly agreed with previous findings (Murphy et al., 2014), including the specific relationship with pupil change and BOLD activity (DiNuzzo et al., 2019).

To develop a more detailed understanding of the temporal relationships between pupil size vs. pupil change with BOLD activity, we performed cross-correlation analyses between both metrics and BOLD. The two pupil features correlated with BOLD on different timescales, with peak correlation between pupil size and BOLD occurring 2.75 s earlier than peak correlation between pupil change and BOLD. Only the correlation with pupil *change* took on the appearance of a typical hemodynamic response, while the correlation with pupil size was already significantly positive at a time-delay of 0 s. This is expected if pupil size is understood as a running integral of pupil change. Functionally, this means pupil size may behave less like a response to any discrete event and more like a slowly varying state variable with which BOLD activity covaries, in a way that does not easily map on to phasic or tonic profiles of LC neurons (Nuiten et al., 2026). Within the LC ROI itself, none of these analyses can adequately resolve whether pupil-BOLD coupling lies within the LC itself or in immediately adjacent tegmentum. What was consistent across both the canonical HRF and cross-correlation analyses was strong pupil change-BOLD coupling within a few millimetres of the LC.

The strength of the peak correlation between pupil metrics and BOLD differed by cortical network, with the strongest association in the unimodal ventral attention network and much weaker correlations in transmodal default mode and limbic networks, consistent with their positions along the principal gradient (Margulies et al., 2016). This profile is consistent with the ventral attention network’s established role in stimulus-driven orienting (Corbetta & Shulman, 2002) and with prior work linking task-evoked pupillary responses to orienting (Riley et al., 2023; Swallow et al., 2022; Wang & Munoz, 2015). Orienting activity can result from either external or internal events (Corbetta et al., 2008). Here we found that the oddball task was a significant, but not sole, source of orienting events, with moments of highest pupil change significantly more likely than chance to occur when oddballs were presented. Considering pupil size as a continuous regressor allowed us to capture both task-driven and task-independent brain and pupillary events.

To better understand the pattern of brain activity associated with pupillary features, we expanded our focus beyond the LC to ask whether they were explained by any of 19 neurotransmitter receptors or transporter maps, drawing on PET datasets made available by Hansen et al., 2022. Pupil size-BOLD coupling on its own was not associated with any PET maps. After partialing out the pupil change-BOLD coupling map, however, unique pupil size-BOLD coupling was explained by μ-opioid (MOR), cannabinoid-1 (CB1) and glutamatergic (mGluR5) receptor density. This analytic step was undertaken to isolate aspects of pupil-BOLD coupling that might be specific to the pupil size vs. pupil change metric, since the two measures are derived from the same continuous signal and their coupling maps shared 36.9% of their spatial variance. Pupil change-BOLD coupling, both originally, and also after pupil size-BOLD correlation was partialled out, was positively correlated only with norepinephrine transporter (NET) density. Pupil change-BOLD coupling was also very strongly anticorrelated with the principal cortical gradient, as was NET density. Notably, NET density had no significant correlation with pupil size-BOLD coupling (r = 0.04 vs. r = 0.42 for pupil change), reinforcing that the two pupil features carry different information. The association between NET density and pupil change-BOLD coupling did not survive removal of the principal cortical gradient, most likely because both NET density and pupil change-BOLD coupling are strongly anticorrelated with that axis. We further found that individual differences in pupillary responses predicted individual differences in pupil change-BOLD association with NET density, even though they were not able to predict LC BOLD responses.

We did not find any association between pupil-BOLD coupling and other neurochemical systems proposed to modulate pupil size: acetylcholine (VAChT, Fotiou et al., 2009), serotonin (several receptors; only negatively associated with pupil change after partialing out the effect of pupil size) (Noehr-Jensen et al., 2009), or dopamine (DAT, Bartošová et al., 2018) . This absence of evidence may not necessarily be readable as evidence of absence. The tracer maps differ substantially in statistical power. In particular, the VAChT map was assembled from a small cohort (n = 18, the only map made from older adults) and the DAT tracer has limited dynamic range in cortex because it was developed for striatal imaging, so null results for those systems should be taken relatively lightly. However, it was still surprising that we did not detect any association between VAChT density and pupil size-BOLD correlation, a result that we were expecting based on Reimer et al., 2016.

The degree to which pupil change coupling was associated with NET density declined with age. This was not true of other significant pupil-BOLD coupling relationships which remained similarly strong within each group. Overall, from these data we conclude that pupillary dynamics represent similar quantities across the adult lifespan, but aspects of pupillary dynamics specifically linked to noradrenergic quantities were modestly but significantly lower in older adults, while pupil size correlations were not, in an apparent dissociation. These differences could not be explained by decreased data quality in older adults; quality was high and comparable across groups, including within the LC. Older adults thus retained NET-aligned pupil change-BOLD coupling and continuous LC-pupil coupling while lacking a detectable oddball-evoked LC response.

The reduced pupil-brain linkage we observed could be the result of reduced activity or norepinephrine signaling from the LC; this hypothesis is consistent with many reports suggesting that LC activity weakens with age (Bell et al., 2023; Chen et al., 2023; Downs et al., 2025; Grudzien et al., 2007; Mather & Harley, 2016). Older adults may have a different spatial pattern of NET expression, which could account for these results. Hansen et al. tested whether age had a significant effect on the reported maps and found that it did not, but this does not rule out age effects not present in the initial data. The NET density map used here was made from 77 individuals with a mean age of 33.4 y (SD = 9.17); older adults were likely not included.

However, the MOR, CB1 and mGluR5 maps were also made from similarly-aged cohorts (i.e., average age in 30s) and we did not observe any significant effect of age in the these effects. Only the VAChT map came from older adults (mean age 66.8 y, SD = 6.8).

The relationship between unique pupil size-BOLD coupling and μ-opioid density was independent of the principal cortical gradient. This relationship was present in all age groups and did not differ between them. μ-opioid receptors are well known for their effect on pupil size (Murray et al., 1983), with exogenous opioids causing pupil constriction. In rodents, there is evidence that endogenous opioid signaling mediates the pupillary light reflex (PLR, Cleymaet et al., 2021). The PLR was modestly tested here, although lighting conditions in the MRI scanner were such that a full PLR could not be elicited. Together with our observation that pupil size behaves more like a slowly varying state variable rather than an event-linked response, these findings raise the possibility that tonic pupil size carries information about opioidergic tone that is distinct from the noradrenergic information carried by pupil change. Because this was the only neurochemical association in our dataset that was independent of the principal cortical gradient, it warrants direct testing in the future.

## Conclusion

Our results suggest that two components of the pupil signal, its size and its rate of change, carry different information about brain activity. These components correlate with brain activity on different timescales, in different locations, and match the spatial distribution of different neurochemical systems. Only the rate of pupil change appears to relate to the norepinephrine system, judged by its correlation with areas of the brain rich in NET. NET marks the terminals of noradrenergic projections which originate from the LC. The LC is likely driving brain-wide activity in moments when the pupil changes size rapidly, and using pupillometry to identify moments of rapid change in pupil size may be a meaningful way to measure noradrenergic activity. These pupillary responses remain significantly and similarly linked to brain activity across the adult lifespan, but a weakening of the relationship may suggest altered LC signaling in older adults. In sum, distinct components of the pupil signal may provide a window into distinct large scale neuromodulatory cortical dynamics, including prominently but not exclusively the LC-NE system.

## Methods

### Participants

We recruited 90 adults aged 19-78 for a study that included structural and functional MRI with concurrent in-scanner pupillometry. The final sample included 89 adults (67 female, 21 male, 1 nonbinary; mean age = 49.7 y, SD = 16.2), and participants were divided into three age groups: younger (< 45 y; N = 35; mean = 32.9, SD = 8.2), middle-aged (45-64 y; N = 33; mean = 54.2, SD = 5.2), and older (≥ 65 y; N = 21; mean = 70.4, SD = 3.3). Of these, 3 did not have pupil data (either due to previous cataract surgery [N = 1], ptosis [N = 1], or equipment failure [N = 1]) but did have MRI data. Conversely, 5 had pupil data but had MRI data collected with a single-echo sequence that was not analyzed due to inadequate quality. One person did not have usable pupil or MRI data. Physiological recordings were unavailable for 12 participants. The data was truncated for 2, failed visual quality control for 3, and was missing for 7, leaving 72 participants for the physiological-noise models. Thus, N=84 for MRI-only analyses, N=86 for pupil-only analyses, N=81 for combined pupil-MRI analyses, and N=72 for analyses with physiological nuisance regressors.

### Study procedures

Participants were screened for diagnosed cognitive impairment, neurological disease, severe head injury, ocular disease, or any MRI contraindication, and had vision and hearing that were normal or correctable to normal. The night before participating, participants were instructed to refrain from drinking alcohol and to get a good night’s sleep if possible. On the day of the study, participants were instructed to consume their usual amount of caffeine but to finish it ≥1 hour before beginning the study. The study took place at Cornell University from October 2022 through December 2025 and was covered under Institutional Review Board protocol 1910009087. The study began at approximately 9:30 AM for all participants, with the MRI given around 1:00 PM.

Before the MRI, participants practiced the oddball task described here outside of the scanner. Once in the scanner, the oddball task was completed approximately 30 minutes after the start of the scan, and 35 minutes before the end of the scan. Immediately before the task began, a 5-point eyetracker calibration and drift check was performed.

### Task details

The oddball task had three types of stimuli: standard (60%), oddball (20%), and bright (20%). The task itself was modeled on the task described by Murphy et al., 2014 to ensure maximum relevance to LC functioning, with the addition of a bright stimulus to elicit a pupillary light reflex, although the required distance from the screen due to the MRI scanner made this effect mild. Standard stimuli were 1.8° visual angle purple circles, while oddball stimuli were 3.6° visual angle purple circles, both presented on a gray isoluminant background. Oddball stimuli required a button press using an MRI-compatible button box. Due to a technical problem with the task, button presses were recorded only for the second half of the sample (44 participants; task performance data is therefore based on these individuals). In the first half, the total number of trials was 180 vs. 300 in the second; the task was revised to increase the pace of stimulus presentation within the same task length to be able to measure oddball responses more accurately. All other details remained identical in both versions. Bright stimuli were the same size and color as standard stimuli but were 48% brighter than standard stimuli. Participants were not told about the bright stimuli. All stimuli were presented for 75 ms each. Between stimuli, participants fixated on a black fixation dot (0.15° visual angle). The inter-stimulus interval was jittered (mean = 2.0 s, median = 1.5 s, range = 0.5-11.0 s). The sequence of stimuli was designed using optseq2 (Fischl et al., 1999). Total task time was 10 minutes.

### Physiological measurement

Heart rate and respiratory activity were continuously recorded at 100 Hz (cardiac) and 25 Hz (respiratory) during the fMRI session. Heart rate was obtained from a photoplethysmography (PPG) sensor placed on the nondominant index finger, and respiration was recorded with a pneumatic respiration belt. These data were used to regress physiological noise as described in *Preprocessing.* Alongside our analyses of the relationships between pupil activity and BOLD activity, we also analyzed the relationships between heart rate and respiration and BOLD activity as a control; see the **Supplement.**

### Pupillometry

Pupil diameter was continuously recorded from the right eye at 1000 Hz with an SR Research EyeLink 1000 Plus during the fMRI session. Raw traces were cleaned using the following procedures: (1) EyeLink blink-flagged samples were removed with a ±50 ms margin; (2) samples for which |Δpupil| > median(|Δ|) + 8×MAD were removed, following (Kret & Sjak-Shie, 2019) and as in our previous work (Riley et al., 2023); (3) gaps ≤ 1500 ms were linearly interpolated while longer gaps were simply not included in group averages.

To measure task-evoked pupillary responses, per-trial traces were baseline-corrected to the 50 ms pre-stimulus window, downsampled to 10 ms, normalized to each participant’s dynamic range (difference between 1st and 99th percentile cleaned pupil size values per participant), averaged within trial type per participant, and combined as group mean ± SE. Dynamic range normalization was used to account for the fact that range decreases across the adult lifespan (Riley et al., 2023).

*Pupil data quality.* Pupil data was divided into trials 2.4s long each. Across all participants, a median of 6.3% of trials were discarded due to inadequate quality (missing more than one third of samples). Among retained trials, an average of 8.4% of samples were missing per trial (SD = 5.6%). We tested age-group differences using separate linear models on the proportion of trials discarded and on the proportion of missing samples per retained trial, each with a fixed effect of age group. Neither the proportion of trials discarded (F[2,78] = 1.38, p = 0.26) nor the proportion of missing samples per retained trial (F[2,78] = 1.19, p = 0.31) differed across age groups.

MRI.

*Acquisition.* We used a 3T GE Discovery MR750 (software DV29.1) with a 32-channel head coil at the Cornell MRI Facility. Participants laid supine on the scanner bed with their head supported and immobilized. Ear plugs and headphones were used to reduce scanner noise. Visual stimuli were presented with a 32” Nordic Neuro Lab liquid crystal display (1920 pixels × 1080 pixels, 60 Hz) located at the head of the scanner bore and viewed through a mirror attached to the head coil.

Anatomical data were acquired with a T1-weighted MPRAGE sequence (TR = 2600 ms; TE = 2.85 ms; TI = 1000 ms; 7° flip angle; phase acceleration factor 2; 1.0 mm isotropic voxels, 200 slices). Multi-echo echo planar imaging (EPI) sequences were used to acquire functional during the task (TR = 2500 ms; TEs = 14, 27, and 40 ms; multiband factor 2; phase acceleration factor 3; 80◦ flip angle; 1.83 x 1.83 x 2 mm voxels; 78 slices). 240 volumes were acquired (600 s), and the first 2 volumes were discarded leaving 238 volumes (595 s) for analysis. To locate the LC, we used a 2D T1-weighted fast spin-echo sequence (ETL = 3, TR = 600 ms, TE = 14 ms, flip angle = 90°, field-of-view 22 x 22 cm, phase FOV = 80%, 6 averages, 0.43 x 0.43 x 3mm resolution with the acquisition plane perpendicular to the brainstem).

*Preprocessing.* Anatomical data were processed with Freesurfer recon-all (Fischl, 2012). This allowed the identification of each participant’s 4th ventricle (for later masking), and the creation of individual gray matter masks. AFNI’s SSWarper2 (P. Taylor et al., 2024) was used to calculate nonlinear warp parameters for each individual to MNI 2 mm standard space.

Multi-echo EPI was processed with AFNI (P. A. Taylor et al., 2018), including despiking, slice-timing correction, EPI-to-anatomical alignment, volume registration to the min-outlier volume (resampled to 2.0 mm isotropic), brain masking, multi-echo combination and denoising with tedana (Community et al., 2023), scaling to percent signal change, and nuisance regression of polynomial drift terms. Motion parameters were not entered as regressors for whole-brain analysis; motion-related variance was instead removed by tedana, and volumes were censored when more than 5% of brain voxels were flagged as outliers (in the pupil-modulated analysis, volumes with framewise displacement greater than 0.5 mm were additionally censored). Voxels in the 4th ventricle (identified using FreeSurfer recon-all on the MPRAGE images) were excluded to reduce any potential for signal contamination in the LC. For the LC ROI analyses, cardiac and respiratory noise was additionally modelled from the physiological recordings: RETROICOR cardiac and respiratory phase regressors (Glover et al., 2000), heart rate convolved with the cardiac response function, and respiration volume per time convolved with the respiration response function, together with the first five principal components of the 4th ventricle signal (Singh et al., 2022).

To locate the LC, we used two different methods. First, and for most analyses, we used FreeSurfer’s Bayesian SegmentAAN algorithm (Olchanyi et al., 2024) and resampled the resulting ROI to each participant’s native EPI grid. As a comparator, we included manually traced ROIs, following our previously published methods (Riley, Cicero, Mabry, et al., 2025), the LC on our specialized fast spin echo anatomical scan. These were only available for 67 of 89 participants (75%). Briefly, two trained raters traced the LC following the algorithm described in (Turker et al., 2021), and voxels included in both tracings were included in the final ROI. If the Dice coefficient was not higher than 0.6, both raters redid the tracing. The average Dice coefficient was 0.69.

In order to determine whether group-level results included significant voxels in the LC, custom probabilistic ROIs for this group of participants were created from both Bayesian and hand-traced LC ROIs by applying the SSWarper2 warp parameters to the ROIs to move them to MNI space. For the hand-traced ROIs, the ROIs additionally needed to be aligned to each individual’s MPRAGE; this was accomplished using AFNI’s *3dAllineate*.

*fMRI data quality.* For each functional scan we quantified temporal signal-to-noise ratio (TSNR), the fraction of volumes censored, the anatomical/EPI alignment (Dice coefficient), and framewise head motion, and tested each for age-group differences (N = 84). Data quality was high and did not differ significantly across age groups within TSNR (younger 83.7, middle 83.4, older 82.8; F[2,81] = 0.03, p = 0.97), censored-volume fraction (younger 4.5%, middle 2.3%, older 4.6%; F[2,81] = 1.93, p = 0.15), anatomical/EPI Dice overlap (0.913, 0.907, 0.908; F[2,81] = 0.85, p = 0.43), or framewise motion (younger 0.10 mm, middle 0.12 mm, older 0.12 mm; F[2,81] = 0.33, p = 0.72).

Within the LC specifically, data quality was not lower in older adults: median LC TSNR was 113, 133 and 141 in younger, middle-aged and older adults (Kruskal-Wallis p = 0.023), the residual SD of the LC response model was 0.85, 0.71 and 0.68 (p = 0.022), the LC label contained a median of 35, 40 and 39.5 1 mm^3^ voxels, and censored volumes (median 4, 3, 4.5, p = 0.53) and framewise motion (0.05, 0.07, 0.13 mm; p = 0.081) did not differ significantly although older adults trended towards more motion.

*General linear modeling of task-evoked BOLD responses.* For each participant, we fit a whole-brain voxelwise general linear model to the preprocessed time series (238 TRs, 595 s) in native space. Each trial type (oddball, standard, and bright) was modeled by convolving its onset times with a 3-function basis set (SPMG3), along with nuisance regressors for scanner drift. Model coefficients were estimated with a generalized-least-squares fit that accounted for the temporal autocorrelation of the fMRI noise (AFNI’s 3dREMLfit), yielding a per-participant response amplitude for each condition at every voxel. These single-subject amplitudes were then carried to the group analyses. Per-participant beta maps were nonlinearly registered to MNI space and entered into a second-level model with fixed effects of task condition, age group, and their interaction, and a random effect of participant. A group gray-matter mask was created by combining participant gray matter masks thresholded at 0.1.

In a second whole-brain model, the oddball response was amplitude-modulated by three trial-by-trial pupil measures, allowing us to investigate where BOLD may covary with pupillary responses. For every oddball trial we extracted three measures from the simultaneously recorded pupil data: (1) the pretrial pupil size, (2) the area under the task-evoked pupil response over the trial, and (3) the peak rate of change (the maximum of the pupil derivative during the trial). Each measure was mean-centered within participant and entered as a parametric modulator of the oddball regressor. All other terms matched the base univariate model.

BOLD responses within the SegmentAAN LC labels were estimated, in native space, for each trial type with a finite impulse response (TENT) model with knots every 2.5 s from −2.5 to 15 s after stimulus onset. Polynomial drift terms, and nuisance regressors as described above (physiological regressors and 4th ventricle components; n = 72 with such data available) were added with outlier volumes censored as in the whole-brain analysis. An additional model with only head-motion and 4th ventricle nuisance regressors was estimated for all 84 participants.

Bilateral LC signals were partial-volume weighted and averaged across hemispheres before model fitting. Curves are displayed from −2.5 to 10 s. Within each age group, each condition’s estimate at each knot from 2.5 to 10 s was tested against zero by a sign-flipping permutation test across participants (10,000 permutations) with the maximum absolute value of t across the three conditions and four knots as the corrected statistic (Nichols & Holmes, 2002). A response was declared only at knots from 2.5 s onward; the −2.5 s and 0 s estimates are shown as checks because a BOLD response at those times is not physiologically possible. To test the specificity of oddball-evoked responses, the same model was applied to control regions (the 4^th^ ventricle, and a shell of pontine tegmentum around the LC), to the SegmentAAN arousal nuclei within 10 mm of the LC (see **Supplement and Supplementary Fig. S4**).

*Continuous, task-naive pupil regressors.* Following Murphy et al. 2014, task-naive pupil regressors (pupil size and pupil first derivative, dP/dt) for each participant were downsampled to TR rate, convolved with AFNI’s GAM HRF and entered directly into a first-level GLM. For group analysis, both regressors and their interaction with age group, plus a random effect of participant, were entered into a second level model.

*Pupil-BOLD cross-correlation.* For each participant, and within each voxel, Pearson correlations were computed between the BOLD time series in native space and two pupil regressors (pupil size and pupil first derivative, dP/dt, in both cases not linked to task trials), in 0.25 s increments between a −2 s to +10 s delay (pupil preceding BOLD). For each delay, the 20 Hz pupil series was shifted and then averaged within each TR window (50 pupil samples within each 2.5 s TR). In order to measure the cross-correlation profile within each of the 7 Yeo networks (Yeo et al., 2011), we applied the reverse of our individual nonlinear native space-to-MNI warps to move Yeo networks from MNI space each participant’s native space, where computation took place. To obtain a single coupling value per voxel per participant, we extracted each participant’s correlation at the fixed delay corresponding to the peak of the group-average cross-correlation profile (+2.25 s for pupil size and +5.00 s for pupil derivative). These coupling (Pearson r) maps were warped to the 2 mm MNI template and entered into a voxelwise one-sample t-test against zero (3dttest++) to identify voxels significantly coupled to pupil size or its derivative across participants. The same procedure was used to compute respiration-BOLD and heart rate-BOLD cross-correlations (**see Supplement and Supplementary Figure S1.**)

*PET tracer analysis methods.* Nineteen receptor/transporter maps covering 9 neurotransmitter systems (acetylcholine, dopamine, endocannabinoid, GABA, glutamate, histamine, norepinephrine, opioid, and serotonin) were drawn largely from Hansen et al., 2022 using the *neuromaps* python package (Markello et al., 2022). Where more than one map was available for the same target, we selected the cohort with the larger sample at 1mm resolution. Our set differs from the atlas as published by Hansen et al. in two respects: we included the κ-opioid receptor ([¹¹C]LY2795050, Vijay et al., 2018) in place of the NMDA receptor ([¹⁸F]GE-179), and we used [¹⁸F]fallypride rather than [¹¹C]FLB-457 for D2/D3 (Jaworska et al., 2020). For each tracer, PET density was averaged within each of the 200 cortical parcels of the Schaefer 200 atlas (z-scored) and correlated (Pearson r) with the parcel-mean group pupil-BOLD (or pupil derivative-BOLD) coupling t-statistic.

*General group analysis principles.* Group-level analyses of pupil data used linear mixed-effects models (*lme4* package in R, Bates et al., 2015) with a random intercept for participant. Group-level analyses of the voxelwise maps used linear mixed-effects models (*3dLMEr* in AFNI), with relevant fixed effects (e.g., trial type, pupil regressor, age group, and their interactions) and a random intercept for participant. Wherever possible, analyses were carried out in each participant’s native space and no spatial smoothing was applied. Individual maps were warped to the 2 mm MNI template only where analysis required a common space. For the whole-brain voxelwise analyses we controlled the family-wise error rate by thresholding voxels at p < 0.001 and retaining only clusters large enough (using a NN=2 definition) to yield a corrected cluster-wise α < 0.05, with the cluster-extent threshold estimated by Monte Carlo simulation of the residual spatial smoothness (AFNI’s 3dClustSim). For analyses relating pupil-BOLD coupling maps (which were calculated in native space) to the PET density maps (after warping the coupling maps to MNI space), spatial correlation was quantified as the Pearson correlation across Schaefer 200 cortical parcels (Schaefer et al., 2018). These were tested against a null distribution of 5,000 rotations (Alexander-Bloch et al., 2018) that preserve each map’s spatial autocorrelation. In all spin tests, the pupil-BOLD coupling map was rotated while the PET density map was held fixed. Across all neurotransmitter systems tested, p-values were corrected with the Benjamini-Hochberg false-discovery-rate procedure (Benjamini & Hochberg, 1995).

## Use of artificial intelligence tools

The initial data analysis code was written by the corresponding author between 2022 and 2026. Beginning in 2026, Anthropic models (Claude; models Opus 5.0/5.5, Fable 5.1) were utilized to optimize code, translate existing MATLAB and R scripts into Python for faster performance and easier sharing, and implement and accurately report new statistical tests (Alexander-Bloch spin tests, permutation-based maximum statistic tests). To proofread the manuscript, the corresponding author used the same models to verify the accuracy of each number reported in the text against the analysis outputs and to populate data tables directly from model results to prevent transcription errors. AI was also used to organize all of the scripts that produce every number reported in this manuscript and place them within a shareable package (see below). AI tools were not used to write or edit the text of the manuscript. The authors take full responsibility for the integrity and content of this work.

## Availability of data and code

All analysis code and de-identified derived data (coupling maps, per-participant coupling and pupillometry measures, and LC region-of-interest estimates) is publicly available at 10.5281/zenodo.22285594.

## Contributions

CRediT authorship contribution statement: **E. Riley:** Conceptualization, Methodology, Software, Validation, Formal analysis, Investigation, Data curation, Writing – original draft, Writing – review & editing, Visualization, Supervision, Funding acquisition, Project administration. **E. De Rosa:** Conceptualization, Funding acquisition, Resources. **A. Anderson:** Conceptualization, Funding acquisition, Resources, Writing – review & editing.

