## Supplementary Fig. S2 for "Pupil size and pupil change correlate with distinct cortical neurochemical systems across the adult lifespan"

### Supplement

#### Supplementary Tables S1. Whole-brain univariate oddball activation (N = 84)

**Main effect of condition (omnibus $\chi^2$) — 99 clusters, 6937 voxels**

| Voxels | x | y | z | Peak $\chi^2$ | Region |
|---|---|---|---|---|---|
| 1635 | -63 | -43 | 39 | 59.54 | L supramarginal |
| 1170 | 57 | -39 | 45 | 47.35 | R supramarginal |
| 962 | 23 | -55 | -17 | 51.81 | R cerebellum / ventral occipitotemporal |
| 449 | -27 | -69 | -23 | 47.62 | L cerebellum |
| 259 | -3 | -65 | 21 | 48.29 | L precuneus / posterior cingulate |
| 223 | 47 | 39 | 21 | 39.56 | R middle frontal |
| 222 | 31 | -47 | -51 | 41.38 | R cerebellum |
| 151 | -47 | -75 | 37 | 31.34 | L inferior parietal |
| 131 | 59 | 11 | 11 | 44.99 | R IFG (opercularis) |
| 113 | -1 | 7 | 37 | 27.28 | middle cingulate |

***The 10 largest clusters are shown (of 99 surviving clusters).***

**Oddball vs standard (Z) — 87 clusters, 5871 voxels (74 oddball > standard, 13 oddball < standard)**

| Voxels | x | y | z | Peak Z | Region |
|---|---|---|---|---|---|
| 1255 | -63 | -43 | 39 | 6.79 | L supramarginal |
| 1219 | 59 | -37 | 45 | 6.63 | R supramarginal |
| 1112 | 23 | -55 | -17 | 6.27 | R cerebellum / ventral occipitotemporal |
| 426 | -29 | -71 | -23 | 6.46 | L cerebellum |
| 246 | 27 | -51 | -53 | 6.05 | R cerebellum |

| | | | | | |
|---|---|---|---|---|---|
| 125 | -3 | -65 | 21 | -5.14 | L precuneus / posterior cingulate (targ<nontarg) |
| 74 | -1 | 27 | 31 | 4.69 | middle / anterior cingulate |
| 70 | 47 | 39 | 21 | 5.15 | R middle frontal |
| 66 | -9 | -87 | 3 | 4.69 | L calcarine |
| 63 | -5 | -43 | 37 | -5.80 | L posterior cingulate / precuneus (targ<nontarg) |

***The 10 largest clusters are shown (of 87 surviving clusters).***

**Oddball: young − old (Z) — 63 clusters, 638 voxels (55 young < old, 8 young > old)**

| Voxels | x | y | z | Peak Z | Region |
|---|---|---|---|---|---|
| 48 | -3 | -17 | 71 | -5.13 | paracentral lobule / SMA |
| 25 | 27 | -51 | -49 | -4.71 | R cerebellum |
| 23 | 5 | -81 | 3 | -5.06 | R calcarine |
| 22 | -11 | -31 | -3 | -4.68 | L thalamus |
| 20 | -5 | -57 | 61 | -4.97 | L precuneus |
| 16 | -57 | -45 | 7 | -4.30 | L middle temporal |
| 16 | 3 | 25 | 29 | -4.93 | anterior / middle cingulate |
| 16 | 9 | -65 | 55 | -5.61 | R precuneus |
| 16 | 43 | -69 | 41 | -5.19 | R inferior parietal |
| 13 | -47 | 3 | -21 | -4.28 | L temporal pole |

***The 10 largest clusters are shown (of 63 surviving clusters).***

**Condition × age interaction (omnibus $\chi^2$) — 2 clusters, 17 voxels**

| Voxels | x | y | z | Peak $\chi^2$ | Region |
|---|---|---|---|---|---|
| 10 | -1 | -19 | 71 | 21.97 | L paracentral |
| 7 | -29 | -85 | 35 | 18.07 | L middle occipital |

**Bright: young − old (Z) — 4 clusters, 34 voxels (all young < old; marginal)**

| Voxels | x | y | z | Peak Z | Region |
|---|---|---|---|---|---|
| **12** | **-29** | **-101** | **13** | **-4.51** | **L lateral occipital** |
| **9** | **21** | **-3** | **-7** | **-4.50** | **R pallidum** |
| **7** | **19** | **-87** | **3** | **-4.04** | **R calcarine** |
| **6** | **19** | **-75** | **47** | **-3.72** | **R precuneus** |

**Null (no surviving clusters): main effect of age group; Bright vs Standard; Standard young−old.**

#### Supplementary Tables S2. Pupil-BOLD coupling (amplitude-modulated GLM)

**Modulator: pretrial pupil size — 736 voxels in 80 clusters (largest 29 voxels)**

| Voxels | x | y | z | Peak Z | Region |
|---|---|---|---|---|---|
| 29 | -13 | -91 | 31 | -4.69 | L superior occipital |
| 27 | 29 | -97 | -5 | -5.90 | R inferior occipital |
| 20 | -15 | -73 | 7 | -4.50 | L calcarine |
| 17 | 3 | -83 | 15 | -5.38 | calcarine |
| 16 | -55 | 1 | 13 | 4.49 | L rolandic operculum |
| 16 | 5 | -93 | 9 | -4.86 | calcarine |
| 14 | 37 | 15 | 41 | 5.20 | R middle frontal |
| 14 | -13 | -79 | 31 | -5.20 | L cuneus |
| 14 | -43 | -57 | 51 | -5.19 | L inferior parietal |
| 12 | 35 | -89 | -5 | -5.32 | R inferior occipital |

*The 10 largest clusters are shown (of 80 surviving clusters).*

**Modulator: pupil derivative — 98 voxels in 13 clusters (largest 11 voxels)**

| Voxels | x | y | z | Peak Z | Region |
|---|---|---|---|---|---|
| 11 | 25 | -99 | 11 | -4.31 | R lateral occipital |
| 11 | 39 | -65 | 11 | -4.74 | R inferior parietal |

| | | | | | |
|---|---|---|---|---|---|
| 9 | 3 | 35 | 25 | 4.41 | R caudal anterior cingulate |
| 9 | -23 | -87 | -37 | -4.12 | L cerebellum |
| 8 | 19 | -79 | 11 | 4.90 | R calcarine |
| 7 | -49 | 1 | -5 | 4.76 | L superior temporal |
| 7 | 41 | 15 | 5 | 4.38 | R pars opercularis |
| 6 | 65 | -3 | 5 | 4.10 | R superior temporal |
| 6 | -1 | 25 | 41 | 3.88 | L superior medial frontal |
| 6 | -3 | -39 | 45 | 3.63 | L precuneus |

*The 10 largest clusters are shown (of 13 surviving clusters).*

**Modulator: pupil AUC — 85 voxels in 10 clusters (largest 13 voxels)**

| Voxels | x | y | z | Peak Z | Region |
|---|---|---|---|---|---|
| 13 | -9 | -93 | 17 | 4.31 | L cuneus |
| 12 | 11 | -83 | 13 | 5.16 | calcarine |
| 9 | -9 | -71 | 7 | 4.50 | L lingual |
| 9 | 45 | -53 | 59 | 4.51 | R superior parietal |
| 8 | -9 | -103 | 21 | 4.33 | L lateral occipital |
| 7 | -15 | -91 | -13 | 3.83 | L lingual |
| 7 | -11 | -71 | 15 | 4.07 | L calcarine |
| 7 | 55 | 9 | 35 | 3.79 | R precentral |
| 7 | -19 | -47 | 59 | -4.47 | L superior parietal |
| 6 | 7 | -73 | 11 | 4.10 | R lingual |

**Omnibus tests ($\chi^2$, one-sided ≥7 voxels)**

| modulator | main-effect cond | main-effect age | cond×age interaction |
|---|---|---|---|

| | | | |
|---|---|---|---|
| baseline | 77 vox (10 clu) | 9 vox (1 clu) | 296 vox (33 clu) |
| derivative | 43 vox (5 clu) | 0 vox (0 clu) | 43 vox (5 clu) |
| AUC | 22 vox (3 clu) | 29 vox (3 clu) | 331 vox (37 clu) |

Supplementary Table S3. Continuous (task-naive) pupil–BOLD coupling

**Regressor: Pupil size X canonical HRF — 214 voxels in 18 clusters (largest 44; 17 negative, 1 positive)**

| Voxels | x | y | z | Peak Z | Region |
|---|---|---|---|---|---|
| 44 | 45 | -23 | 59 | -4.30 | R precentral gyrus |
| 24 | 1 | 57 | -11 | -4.88 | L mid orbital gyrus |
| 20 | 45 | -69 | 1 | -4.68 | R middle temporal gyrus |
| 18 | -13 | -95 | 27 | -4.28 | L superior occipital gyrus |
| 17 | -21 | -91 | 21 | -4.68 | L superior occipital gyrus |
| 10 | -39 | -73 | -9 | -3.91 | L fusiform |
| 8 | 33 | -51 | -33 | 4.52 | R cerebellum |
| 8 | 27 | -95 | 15 | -4.17 | R superior occipital gyrus |
| 8 | 1 | -29 | 63 | -4.30 | R paracentral lobule |
| 7 | 63 | -17 | -1 | -3.85 | R superior temporal gyrus |

*The 10 largest clusters are shown (of 18 surviving clusters).*

**Regressor: Pupil derivative X canonical HRF — 24,948 voxels in 132 clusters (largest 22,087; 103 positive, 29 negative)**

| Voxels | x | y | z | Peak Z | Region |
|---|---|---|---|---|---|
| 22087 | -15 | -71 | 5 | 9.26 | L lingual (contiguous; spans occ/par/insula/striatum/cerebellum/brainstem) |
| 287 | 31 | 47 | 29 | 6.58 | R rostral middle frontal |

| | | | | | |
|---|---|---|---|---|---|
| 286 | -41 | 35 | 29 | 6.60 | L rostral middle frontal |
| 169 | -51 | -71 | 43 | -6.19 | L inferior parietal |
| 148 | 31 | -55 | -59 | 5.76 | R cerebellum |
| 103 | 39 | 3 | -11 | 5.65 | R insula |
| 89 | 3 | -77 | -41 | 6.97 | R cerebellum |
| 83 | -19 | -95 | -11 | 5.27 | L inferior occipital gyrus |
| 82 | 5 | 7 | -3 | 5.14 | R caudate |
| 81 | -1 | -33 | -39 | 5.90 | brainstem |

*The 10 largest clusters are shown (of 132 surviving clusters).*

#### Preprocessing of physiological data (respiration, heart rate, and heart-rate variability).

Respiration (respiration belt, 25 Hz) and cardiac pulse (PPG, 100 Hz) were recorded continuously during the oddball task fMRI scan. For each participant we discarded the first 35 s of each recording (matching removal of the first two fMRI volumes, plus 30 s recording before the onset of each scan). Instantaneous respiratory rate was derived from the respiration belt using *neurokit2* (Makowski et al., 2021), which detects individual breaths and returns respiratory rates. Instantaneous heart rate was derived from the PPG signal, also using *neurokit2*, by identifying systolic peaks with beat-artifact correction. The raw data was upsampled 4x before peak detection using Fourier interpolation. For heart rate variability (HRV), we computed a continuous HRV time series as the root mean square of successive RR differences (RMSSD) within a 30 s window, advanced one beat at a time. After identifying instantaneous heart rate, HRV, and respiratory rate, each series was resampled to 20 Hz to match the sampling rate of the final pupil pipeline, and then z-scored. Recordings were screened for physiological plausibility: cardiac RR intervals shorter than 0.4 s or longer than 2.0 s (less than 30 or more than 150 bpm) and breath-to-breath intervals shorter than 2 s or longer than 15 s were flagged as artifacts. Final quality control for heart rate was performed visually using Poincaré plots. Eight participants were excluded at this step for insufficient data quality, leaving usable heart-rate, HRV and respiratory-rate time series for 73 participants.

#### Physiological and susceptibility/dropout controls.

To confirm that the pupil-BOLD coupling and its neurochemical associations were not driven by physiological confounds or by regional signal dropout, we ran two controls. First, we cross-correlated heart rate, HRV (RMSSD) and respiration rate with the mean whole-brain BOLD signal across delays of −2 to +10 s, exactly as for the pupil measures (Fig. S1 A-D; N =

73 participants with usable physiological recordings). Heart rate was positively correlated with whole-brain BOLD, peaking at $r = +0.234$ (SE = 0.018) with BOLD following heart rate by 2.00 s. Respiration rate showed a weaker and broader positive correlation, peaking at $r = +0.124$ (SE = 0.019) at 5.25 s. HRV showed no peak ($|r| \leq 0.033$ at all delays). For comparison, in the same participants the peak correlation with whole-brain BOLD was stronger for heart rate than for pupil derivative (paired $t[72] = 6.02$, $p < 0.001$), and did not differ between respiration rate and pupil derivative ($t[72] = 0.42$, $p = 0.67$).

We then computed fixed-delay voxelwise maps for each physiological signal at its peak delay and thresholded the group maps at the same cutoff used for pupil-BOLD coupling ($p < 0.001$, cluster ≥ 16). Heart rate was significantly correlated with BOLD in 85,675 voxels, all positive, nearly all within a single cluster covering 27% of brain voxels (peak $t = 10.82$ at MNI [−21, −49, −17]). Respiration rate was significantly correlated with BOLD in 16,410 voxels, all positive, in 51 clusters, the largest in right angular gyrus, and right inferior lateral occipital cortex. HRV yielded no significant voxels. By comparison we found 14,078 significantly correlated voxels for pupil size and 33,629 for pupil derivative. Because heart rate and respiration were themselves coupled with BOLD, we tested directly whether they accounted for pupil-BOLD coupling. For each participant we regressed heart rate, respiration rate and HRV, each at delays of 0, 2.5, 5.0 and 7.5 s (to avoid assuming a particular vascular response latency), out of both the BOLD time series and the pupil regressors, and recomputed the voxelwise pupil size (+2.25 s) and pupil derivative (+5.00 s) coupling maps. Without this correction, group maps from these 73 participants matched the full-sample maps (spatial $r = 0.972$ for pupil size and 0.979 for pupil derivative). Removing physiological signals reduced coupling strength (mean r within significant voxels: 0.034 to 0.030 for pupil size; 0.029 to 0.022 for pupil derivative), but the spatial pattern of coupling was preserved (spatial correlation of group t-maps before vs. after correction: $r = 0.945$ for pupil size, $r = 0.958$ for pupil derivative), and 99% of voxels that remained significant for pupil derivative lay within the uncorrected map. The significant neurochemical associations were unchanged: after correction, pupil derivative coupling remained correlated with NET density ($r = +0.39$, spin $p = 0.003$, vs. $r = +0.42$, $p = 0.001$ without correction), as did pupil derivative coupling with pupil size partialled out (NET: $r = +0.47$, $p = 0.005$; MOR: $r = -0.57$, $p = 0.008$; CB1: $r = -0.52$, $p = 0.012$), and pupil size coupling with pupil derivative partialled out remained correlated with MOR density ($r = +0.58$, $p < 0.001$). Thus pupil-BOLD coupling and its neurochemical associations were not attributable to cardiac or respiratory fluctuations.

We note that the heart rate coupling map, which is also expected to represent orienting responses, was itself spatially correlated with NET ($r = +0.38$, spin $p = 0.032$), MOR ($r = -0.58$, $p = 0.018$) and CB1 density ($r = -0.47$, $p = 0.032$).

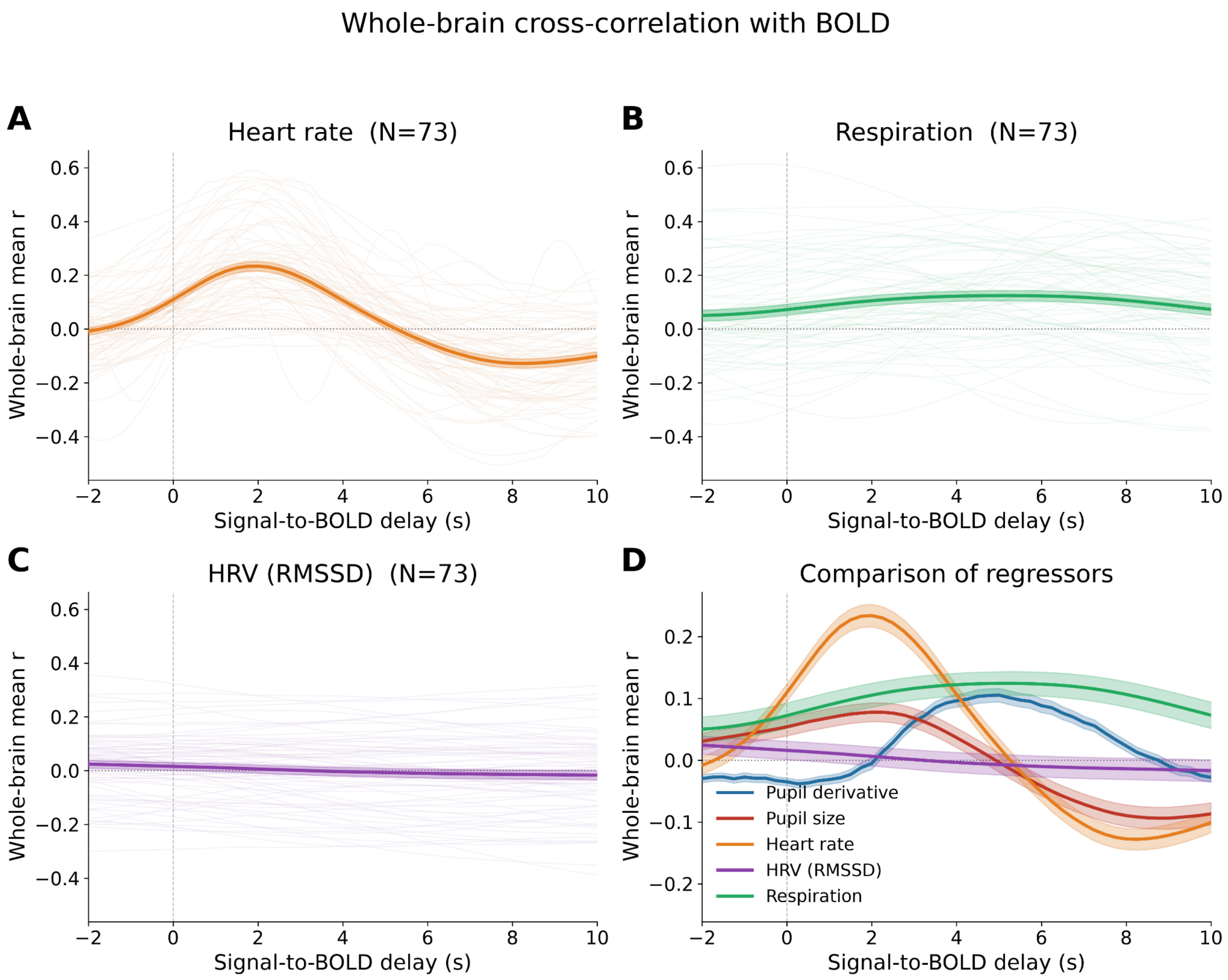


**Fig. S1.** Heart rate-BOLD (A), respiration-BOLD (B), heart rate variability-BOLD (C) and for comparison, all modality (D) cross-correlations across time delays -2 to +10 s. Pearson r between each modality and mean BOLD signal in the whole brain as a function of time delay. Individual traces shown for A-C, shaded bands show ± SE.

Within the LC, we examined the effect of successive nuisance models on the LC oddball response in younger adults (**Fig. S2**). Adding physiological nuisance regressors (RETROICOR cardiac and respiratory phase regressors, heart rate convolved with the cardiac response function, and respiration volume per time convolved with the respiration response function) to the LC model increased, rather than decreased, the oddball response at 2.5 s. The response was not significantly changed by the further addition of fourth-ventricle components or head-motion regressors. We also used the same set of physiological nuisance regressors to investigate modelled LC impulse responses using the SPMG3 basis function (**Fig. S3**).

To test whether putative LC responses were actually specific to the LC, we also examined oddball-evoked responses in the SegmentAAN nuclei (Olchanyi et al., 2024) with centroids within 10 mm of the LC (parabrachial complex, laterodorsal tegmental nucleus, oral pontine reticular formation, median and dorsal raphe), and in two control regions (a 1-4 mm shell of pontine tegmentum surrounding the LC and not sharing any full or partial voxels with the LC or 4th ventricle, and the 4th ventricle itself), by age group, using the same model as in the previous paragraph (**Fig. S4**). We found that the LC had the strongest oddball-evoked response, but significant responses were also observed within the parabrachial complex, oral pontine reticular formation, and the shell of pontine tegmentum. The former two findings are not unexpected given the role of these regions in salience detection and orienting; the latter finding could be caused by registration challenges, hemodynamic spread, or simply true oddball-evoked activity in this area.

We also repeated the cross-correlation analysis described in Section 7 of the main text within the LC ROI. As before, we computed the cross-correlation between the continuous pupil and pupil derivative time series (each separately) and mean LC BOLD across pupil-to-BOLD delays of -2 to +10 s (step size 0.25 s, **Fig. S5**). At the fixed delays used in the cortical analyses, LC BOLD correlated positively with pupil size at +2.25 s (mean $r = 0.033$, $t(71) = 4.57$, $p < .001$) and with the pupil derivative at +5.00 s (mean $r = 0.019$, $t(71) = 2.32$, $p = .023$), N = 72, physiological and 4th-ventricle regressors removed.

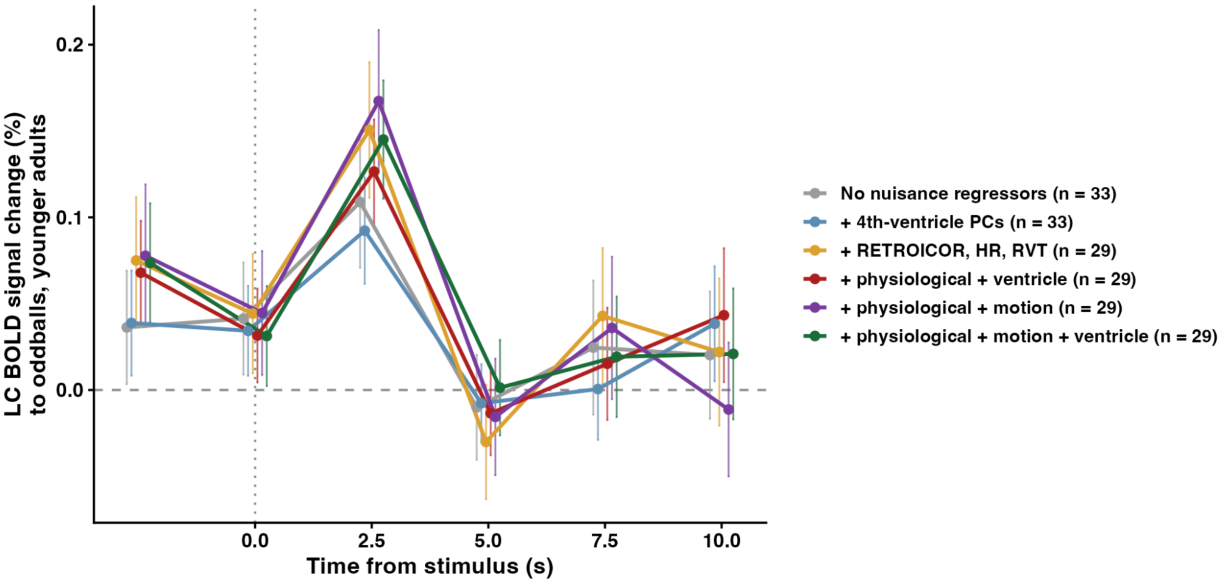


**Fig. S2.** LC oddball response in younger adults under successive nuisance models. Group mean ± SE finite impulse response estimates of the LC BOLD response to oddball stimuli in younger adults, from the model with polynomial drift terms only (n = 33) through the addition of fourth-ventricle principal components, cardiac and respiratory regressors (RETROICOR, heart rate [HR], respiration volume per time [RVT]), and head-motion parameters (n = 29 for models with physiological regressors). Points are offset slightly along the time axis for legibility.

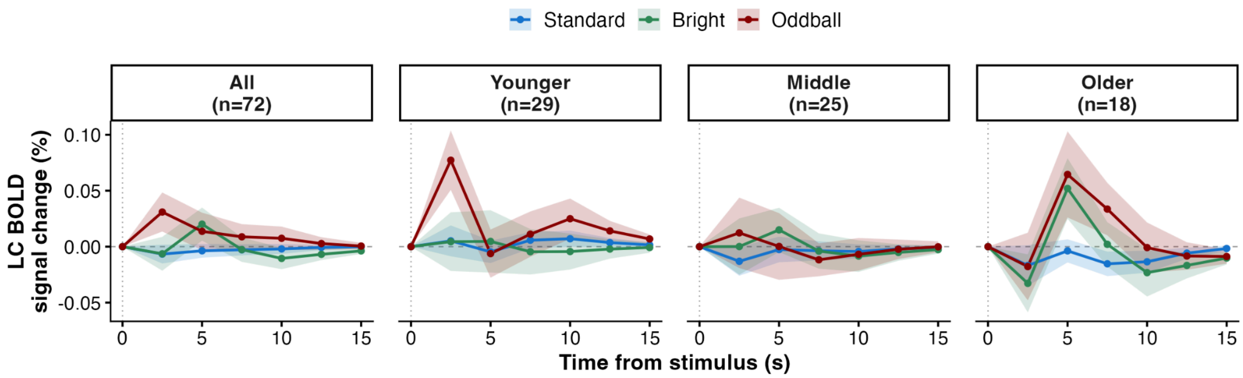


**Fig. S3.** Modelled LC impulse response with physiological noise regressors. For comparison with the finite impulse response estimates in Fig. 1B, the LC response estimated with the SPMG3 basis set (canonical hemodynamic response and its temporal and dispersion derivatives) under the same nuisance model ($n = 72$), by trial type and age group. The fitted response of younger adults to oddballs exceeds that to standards at 2.5 s ($M = 0.072\%$, $t(28) = 2.57$, $p = .016$), so the finding does not depend on the choice of basis.

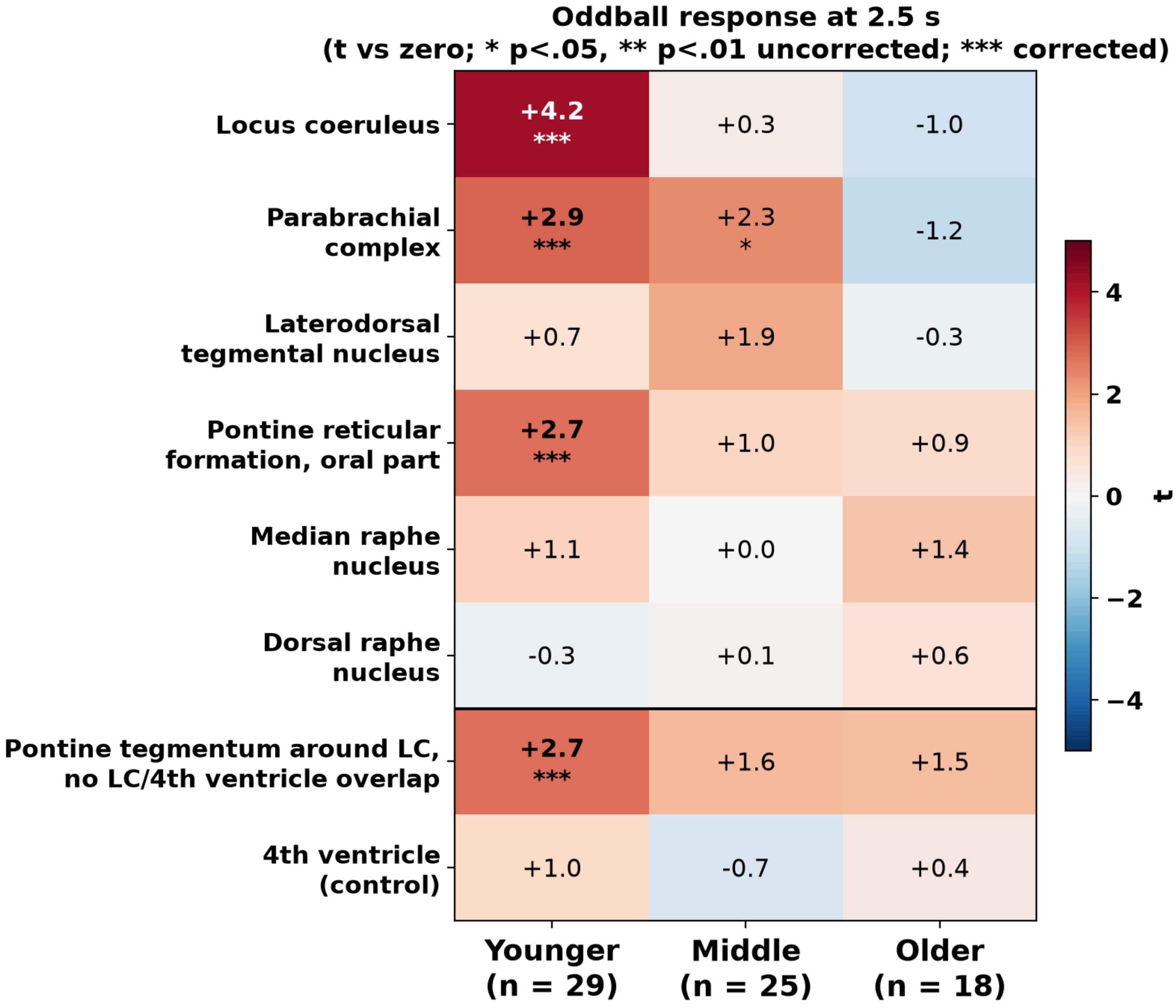


**Fig. S4.** Spatial specificity of the LC oddball response. Group t statistics (response vs. zero) at 2.5 s after stimulus onset for the oddball condition in the SegmentAAN nuclei (Olchanyi et al., 2024) whose centroid lies within 10 mm of the LC (parabrachial complex, laterodorsal tegmental nucleus, oral pontine reticular formation, median and dorsal raphe), and in two control regions (shell of pontine tegmentum surrounding the LC, 4th ventricle), by age group, from the same model as Fig. 1B (n = 29, 25, 18). * $p < .05$ and ** $p < .01$ uncorrected; *** corrected across knots 2.5-10 s within each region by the permutation maximum-statistic procedure with no correction across regions. Bilateral nuclei merged.

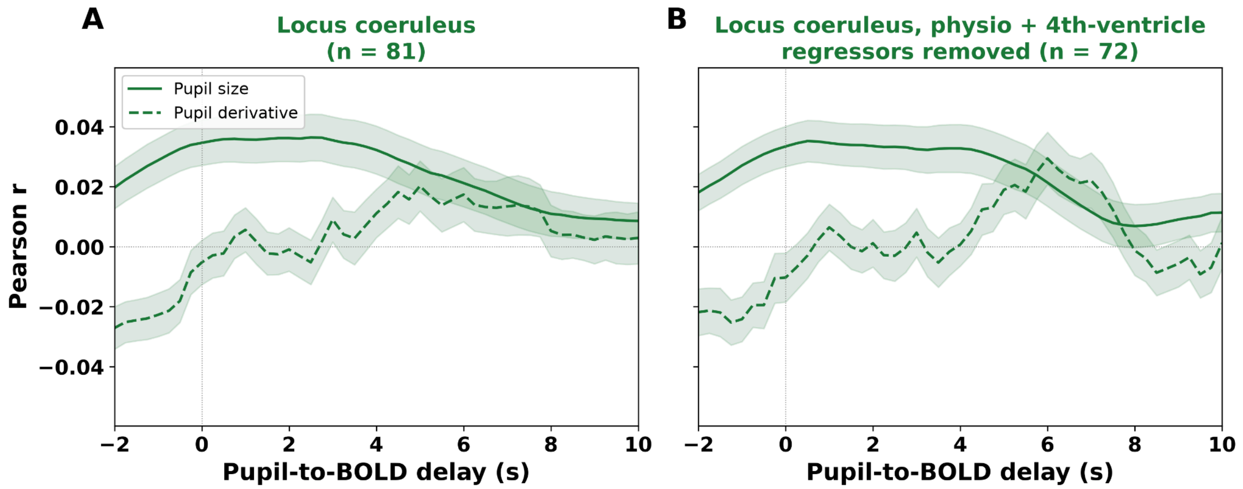


**Fig. S5.** Pupil-BOLD cross-correlation across time delays between -2 to +10 s within the LC. Cross-correlation (r) between pupil and mean BOLD signal as a function of time delay. (A) without physiological regressors and (B) with physiological regressors. Solid lines = pupil size; dashed = pupil derivative. Shaded bands show ± SE.

To test whether the cortical findings presented in the main text depend on physiological noise, we recomputed the parcel-wise pupil-BOLD coupling maps after regressing head motion and fourth-ventricle principal components (n = 81), physiological regressors and fourth-ventricle components (n = 72), or all three (n = 72) from each participant's parcel time series. The group pupil derivative coupling map was spatially correlated with NET density at r = +0.46 in the main model and r = +0.47, +0.45 and +0.47 under the three nuisance models (all spin $p < .001$).

Last, we tested whether the pupil-BOLD coupling maps reflected susceptibility/dropout artifacts, using a group-average T2* map from the same 81 participants (*tedana*) as a reference. We tested whether significant neurochemical PET map correlations with pupil-BOLD coupling maps survived partialling T2* out of the coupling maps (Table S4). Two features of this test should be noted. First, the T2* map is itself spatially correlated with NET density (r = +0.42, spin p = 0.025), so partialling T2* out of any coupling map also removes variance that the coupling map shares with NET; the test is therefore conservative for NET in particular. Second, the residualized coupling maps showed no significant spatial alignment with the T2* map (pupil size coupling with derivative removed, r = +0.05, p = 0.77; pupil derivative coupling with size removed, r = +0.30, p = 0.24). With that caveat, the NET correlation with the raw pupil derivative-BOLD coupling map was attenuated to trend level after T2* removal (r = +0.27, p = 0.078), whereas the NET correlation with the residualized derivative map remained significant (r = +0.39, p = 0.018). The MOR and CB1 correlations were unaffected. The correspondence between pupil-BOLD coupling and PET tracer density is therefore not explained by dropout artifacts.

Table S4. Receptor associations before and after removing T2* map (spin test, N=5000).

| **Finding** | **r (original)** | **r (T2* removed)** |
|---|---|---|
| NET density correlation with pupil derivative-BOLD coupling | +0.42 (p=.001) | +0.27 (p=.078) |
| NET density correlation with pupil derivative-BOLD coupling (size coupling partialled out) | +0.50 (p=.004) | +0.39 (p=.018) |
| MOR density correlation with pupil size-BOLD coupling (derivative coupling partialled out) | +0.58 (p=.0002) | +0.58 (p=.0002) |
| CB1 density correlation with pupil size-BOLD coupling (derivative coupling partialled out) | +0.43 (p=.002) | +0.44 (p=.002) |
